# A CYFIP1-Inspired Peptidomimetic Modulates eIF4E-Dependent Translational Control in Cancer and Neurodevelopmental Disorders

**DOI:** 10.64898/2026.05.06.722988

**Authors:** Alice Romagnoli, Omar Alsina, Stefano Raniolo, Alessandro Gori, Kyriaki Foka, Anastasia De Luca, Jacopo Sgrignani, Jesmina Rexha, Agnese Roscioni, Greta Bergamaschi, Zaira Boussadia, Rita Pepponi, Gloria Venturini, Andrea Cavalli, Tiziana Borsello, Alberto Martire, Anna La Teana, Claudia Bagni, Daniele Di Marino, Vittorio Limongelli

## Abstract

The eukaryotic translation initiation factor 4E (eIF4E) is a central regulator of cap-dependent translation and a compelling pharmacological target in disorders marked by protein synthesis dysregulation, including cancer and Fragile X Syndrome (FXS). Among endogenous eIF4E regulators, the CYFIP1-eIF4E interaction is uniquely selective, offering a framework for designing targeted translation modulators. Here, we report Cy-9B, a rationally engineered, stapled peptidomimetic derived from CYFIP1 that binds eIF4E, disrupts eIF4E-eIF4G complex, and suppresses cap-dependent translation.

Enhanced-sampling free-energy simulations reveal that Cy-9B engages eIF4E through a non-canonical binding mode. Cy-9B exhibits drug-like properties, including high proteolytic stability and nanomolar affinity. Functionally, Cy-9B inhibits lung cancer cell proliferation, migration, and invasion. In neurodevelopmental disease models, Cy-9B partially normalizes excessive translation in FXS hippocampal neurons and rescues social behavior deficits in a Cyfip1 haploinsufficient *Drosophila melanogaster* model, restoring wild-type–like performance. Cy-9B emerges as a first-in-class therapeutic candidate for disorders sharing translational dysregulation, highlighting targeted modulation of eIF4E as a broadly applicable and physiologically compatible therapeutic strategy.

## Introduction

Cancer and Fragile X Syndrome (FXS) are distinct pathological conditions sharing an intriguing molecular overlap that underscores a common dysregulation in protein synthesis. Both disorders, despite their diverse manifestations and severity, converge on impaired protein production, a process vital for maintaining cellular homeostasis and critical functions, including synaptic plasticity and memory in the brain ^1,2^. Notably, this convergence centers on dysfunction in cap-dependent translation, a fundamental mechanism that governs the initiation of protein synthesis and underpins numerous cellular and neurological processes. Cap-dependent translation is initiated when the eukaryotic Initiation Factor 4E (eIF4E) recognizes the m^7^GTP cap at the 5′ end of the mRNA. This recognition facilitates the assembly of the eIF4F complex, which comprises eIF4E, the DEAD-box RNA helicases eIF4A, and the scaffold protein eIF4G ^3^. Assembly of the complex is a rate-limiting step that aids in positioning the ribosome at the mRNA’s start codon to initiate protein synthesis ^4^. eIF4E is one of the least abundant translation factors and thus its expression, activity, and availability are subjected to fine-tuned regulation ^3,5^. In both cancer and FXS, this dysregulation precipitates the aberrant synthesis of specific proteins that contribute to cell malignant transformation and structural anomalies of dendritic spines, respectively. In cancer, uncontrolled cell growth results from the unrestrained synthesis of proteins pivotal for cell cycle progression, survival, and evasion of cellular checkpoints. Accordingly, cancer cells frequently show elevated activity of eIF4E that has been linked to growth stimulation, oncogenic proliferation, and metastasis by selectively enhancing the translation of long and highly structured 5’UTR mRNAs ^6^. These transcripts are tumor-promoting factors (*e.g.,* vascular endothelial growth factor VEGF, cyclins D1, c-MYC, etc.), whose unbalanced translation drives tumor development and progression ^2,7^. The dysregulation of eIF4E is commonly observed in different cancer types, including colon, breast, and lung, in which aggressive tumor behavior and worse clinical outcomes have been correlated with high eIF4E expression ^8,9^. Lung cancer is the second most common cancer and the leading cause of cancer-related death globally, due to late-stage diagnosis and the aggressive nature of the disease ^10^.

Conversely, FXS, the most common inherited cognitive developmental disability, exhibits disrupted synaptic plasticity attributed to abnormal protein synthesis within neurons. The molecular cause of FXS is the expansion of a trinucleotide repeat sequence within the *FMR1* gene which causes the absence of the Fragile X Messenger Ribonucleoprotein 1 (FMRP), an RNA-binding protein crucial for the control of expression of neuronal mRNAs involved in brain development and spine morphology ^11^. Currently, there is no cure for FXS, and the therapeutic interventions aim at managing symptoms and improving quality of life ^12^.

Remarkably, both scenarios implicate the dysregulation of cap-dependent translation as the culprit behind these perturbations. The identification of this shared mechanism suggests novel therapeutic avenues. Strategies aimed at modulating cap-dependent translation, particularly by targeting the cap-binding protein eIF4E and its interactions, hold promise for addressing the intricate challenges posed by cancer and FXS ^13–17^. The eIF4E interaction with the 4E-binding proteins (4E-BPs), a group of proteins that physically sequester eIF4E from eIF4G in a mutually exclusive interaction, is one of the most significant negative controls on eIF4E. In fact, 4E-BPs and eIF4G share a conserved 4E-binding motif (*i.e.*, YXXXXXLΦ, where X: variable and Φ: hydrophobic) that adopts an α-helical conformation to specifically recognize a region in the dorsal surface of eIF4E ^5,18–22^. Among the 4E-BPs, the Cytoplasmic FMRP Interacting Protein 1 (CYFIP1) is a moonlighting and versatile protein involved in two distinct pathways and characterized by a conformational butterfly-like motion ^20,23,24^. When assuming a globular tertiary structure, CYFIP1 functions as a 4E-BP, forming a complex with FMRP and eIF4E, thus preventing the translation of some important mRNAs. Therefore, the absence of FMRP causes the FXS synaptopathy, which is characterized by increased neuronal mRNA translation and deficits in the structure and plasticity of synapses ^25–27^. Conversely, when CYFIP1 has a planar conformation, it forms a hetero-pentameric complex with various other partners, constituting the WAVE Regulatory Complex (WRC) that regulates actin polymerization, essential for processes like cell migration, adhesion, and shape changes ^24,28,29^. A healthy balance between these two functions is essential for neuronal architecture and functionality in a physiological state.

CYFIP1 has been linked to a spectrum of neurodevelopmental and neuropsychiatric disorders, including FXS, autism spectrum disorder (ASD), schizophrenia, and epilepsy ^23,30^. Previous works show that CYFIP1 haploinsufficiency produces domain-specific cognitive deficits and behavioral abnormalities across species, *i.e.*, humans, rodents, and flies ^31–33^. Notably, heterozygous loss of the Drosophila CYFIP ortholog (*Cyfip*^85.1^*/+*) provides a tractable platform for quantifying social behaviors and recapitulates ASD- and schizophrenia-relevant phenotypes observed in broader neurodevelopmental disorder (NDD) contexts^32,34^.

Furthermore, emerging data suggest that CYFIP1 is involved in cancer invasion and metastasis, even if the precise mechanism by which CYFIP1 may contribute to the onset and progression of cancer remains to be explored ^35,36^. Because of its ubiquitous nature, protein synthesis has long been considered a challenging target for therapeutic intervention. However, recent advances in understanding the structural and functional dynamics of translation factors have unveiled new opportunities for selective and precise modulation. Numerous eIF4E structures have been resolved, both in their apo form and in complexes with fragments of their binding partners, such as eIF4G and 4E-BPs. Leveraging these structural insights, several peptides and small molecules targeting eIF4E have been developed ^37–45^. These findings provided the rationale for designing a novel CYFIP1-derived peptidomimetic. Structurally, CYFIP1 is considered an unconventional 4E-BP due to its longer and more degenerate eIF4E-binding sequence (*i.e.*, LDKRLRSECK), which forms a distinctive amino acid interaction pattern with eIF4E ^21–23^. This unique binding interaction presents a valuable opportunity to enhance target specificity in drug design. In fact, the variation in CYFIP1 arrangement provides a more flexible and modular scaffold for structure-based peptides design and optimization. Additionally, CYFIP1 is the only 4E-BP with a full-length PDB structure. The precise localization of its α-helical eIF4E-binding region further informed the design process ^28,46^.

Building on these insights, we developed and optimized a CYFIP1-derived peptidomimetic, named Cy-9B, incorporating a specific chemical modification (*i.e.*, staple) to stabilize its helical conformation. Peptides and peptidomimetics are an emerging class of protein-protein interaction (PPI) inhibitors ^47–49^. Among this class, stapled peptides are known for their ability to enhance peptide stability, bioavailability, and specificity ^40,50^. In recent years, several stapled peptides have shown remarkable potential for treating various diseases, including cancer ^12,51,52^. However, this approach has not yet been applied to FXS, making Cy-9B a pioneering candidate in this context.

Using an interdisciplinary strategy that integrates state-of-the-art molecular simulations with biochemical and cellular experiments, we show that Cy-9B binds eIF4E with nanomolar affinity, disrupts the eIF4E-eIF4G initiation complex, and markedly suppresses cap-dependent protein synthesis. We validate the therapeutic relevance of Cy-9B across complementary disease models spanning oncology and neurodevelopment, including lung cancer cells, FXS hippocampal neurons, and an *in vivo* Cyfip1-haploinsufficient *Drosophila melanogaster* model, underscoring both robustness and translational impact. Importantly, Cy-9B engages eIF4E through a binding mode that is distinct from canonical 4E-BPs, providing a mechanistic framework for structure-guided optimization of selective translation modulators. Together, our results position Cy-9B as a first-in-class CYFIP1-inspired peptidomimetic and establish targeted inhibition of cap-dependent translation as a broadly actionable strategy for cancers and neurodevelopmental disorders characterized by dysregulated protein synthesis.

## Results

### Design and binding free energy evaluation of CYFIP1-derived peptides targeting eIF4E

CYFIP1 is a 4E-BP characterized by the presence of a peculiar variant of the canonical 4E-binding motif folded in a short α-helix (Fig. S1). This variation leads to the formation of a unique and distinct amino acid interaction pattern with eIF4E ^21^. This short α-helix inspired the design of CYFIP1-derived peptides (Fig. 1A) whose binding mechanism to eIF4E was studied using funnel-metadynamics (FM) free energy calculations. First, we evaluated the propensity of the CYFIP1-derived peptide to maintain the native α-helix conformation in solution, building three simulative systems: i) CYFIP1 linear peptide (*i.e.*, Cylin); ii) Cy-1B, where a triazole-bridged macrocyclic scaffold (*i.e.*, staple) was added at the N-terminus of the peptide; iii) Cy-9B, where the staple was added at the C-terminus (Fig. 1B-C). 500 ns of MD simulation of each peptide (*i.e.*, Cylin, Cy-1B, and Cy-9B) in solution was performed to evaluate the stability of the α-helix, which represents the bioactive conformation of the peptide (Fig. 1D and Video S1). By looking at the 4E-binding motif in the CYFIP1 X-ray structure, it was calculated that 60% of the 14 amino acids composing the motif are in α-helix and the remaining 40% are in random coil (Fig. 1D). Comparing the structures of the three peptides sampled along the simulations with the X-ray structure it was clear that the only one that resembles the native conformation is peptide Cy-9B (Fig. 1D and Video S1). Indeed, on average, 58% of the Cy-9B stapled peptide residues are in α-helix (Fig. 1D). These data prompted us to select Cy-9B for the following binding study to eIF4E using FM calculations (Fig. 2) ^53–61^. In FM, the molecular binding process is investigated by enhanced sampling simulations where a bias potential is adaptively constructed as a sum of Gaussian functions in the space of the system’s selected degrees of freedom, namely collective variables (CVs). During the simulation, a funnel-shaped restraint potential is applied to the target system, in our case, eIF4E. This potential is a combination of a cone restraint that includes the binding site and a cylindrical part that is directed towards the solvent (Fig. 2A). As the other partner, Cy-9B, reaches the edge of the funnel, a repulsive potential is applied to the system, disfavouring it from visiting regions outside the funnel. In doing so, the method enables multiple binding events and the exploration of long timescales (from ms to s), while also accounting for the full conformational flexibility of the proteins and solvent molecules. At the end of the calculation, the binding free energy (bFES) is computed, and the binding mode is identified as the lowest energy minimum. FM has been successfully used to investigate complex molecular binding processes ^62–67^.

**Fig. 1.**
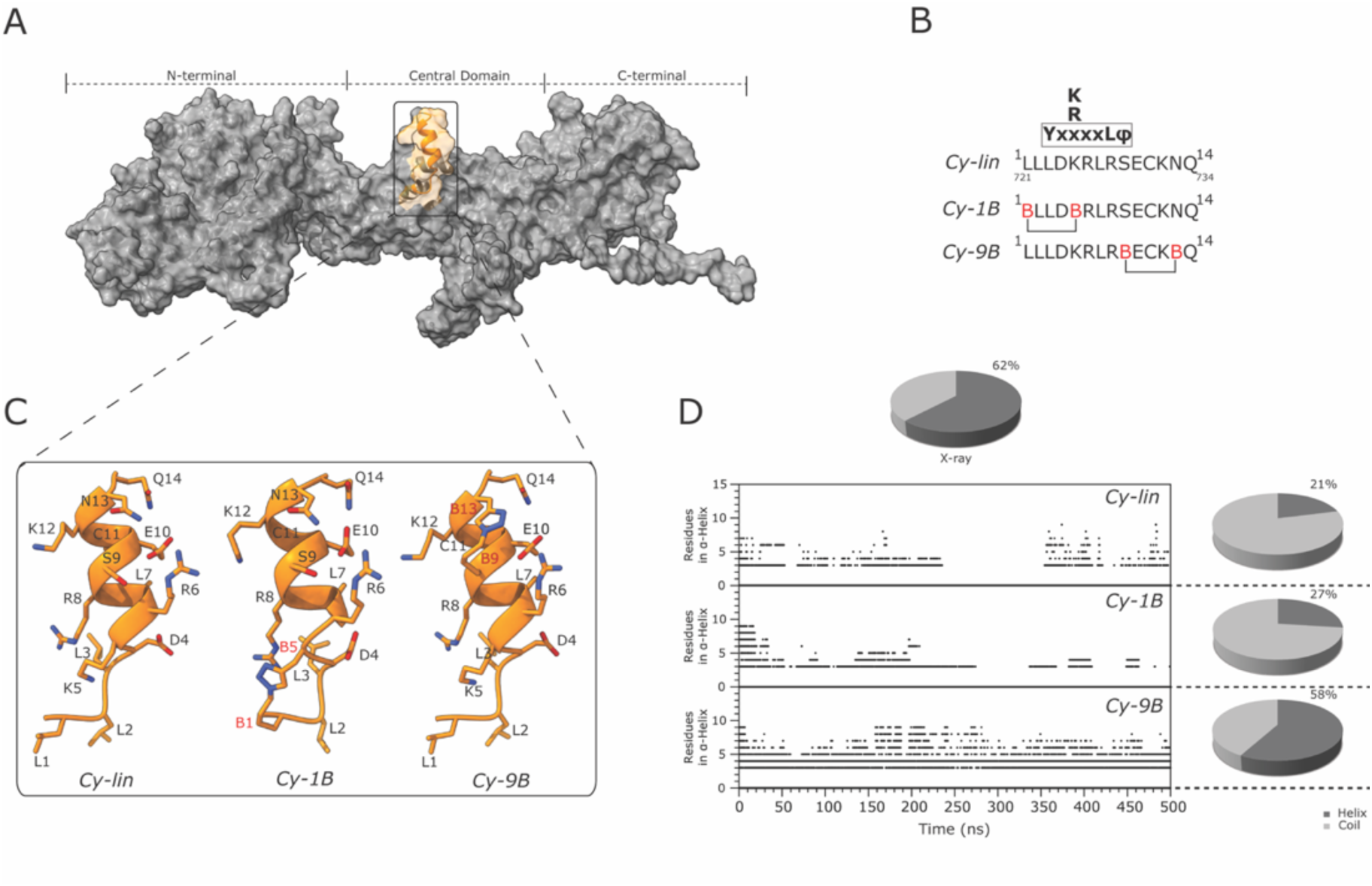
*In silico* design of Cy-9B. **(A)** Three-dimensional structure of CYFIP1, with the 4E-binding motif highlighted in orange. **(B)** Sequence of the canonical 4E-binding motif (bold), together with the linear peptide (Cylin) and the two stapled derivatives (Cy-1B and Cy-9B). The positions of chemical bridges are indicated by the letter B. **(C)** Ribbon representation of the Cylin, Cy-1B, and Cy-9B peptide structure. **(D)** Number of residues adopting an α-helical conformation during molecular dynamics simulations for Cylin, Cy-1B, and Cy-9B. The corresponding average values and those derived from the CYFIP1 X-ray structure (PDB ID: 3P8C) are shown. The graphs next to each plot indicate the average percentage of α-helical residues over 500 ns of simulation, whereas the α-helical content observed in the experimental structure is reported on top.

**Fig. 2.**
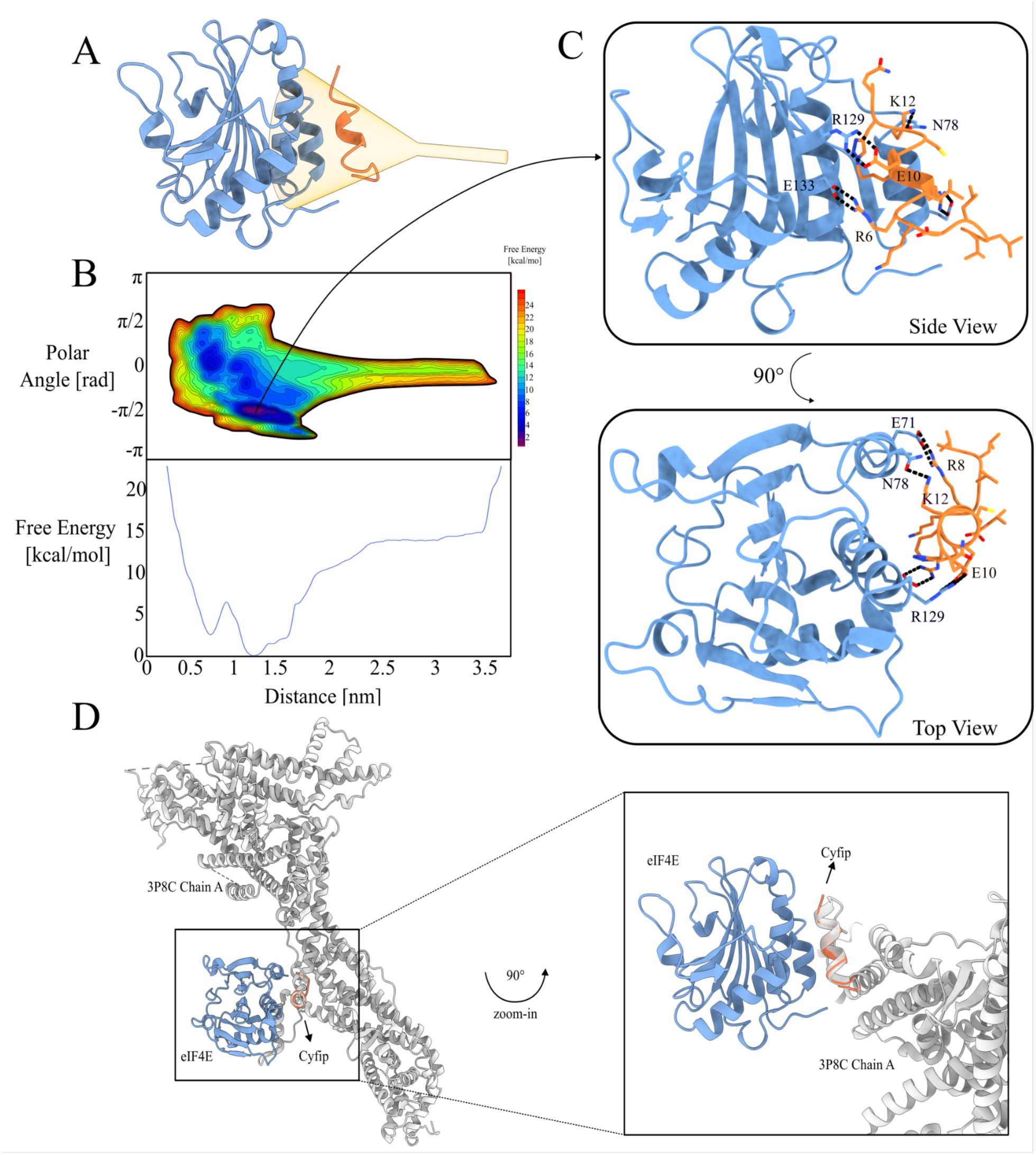
Binding mode of Cy-9B to eIF4E. **(A)** Representation of the funnel used in the simulation system. The target protein eIF4E is shown as a blue cartoon, and the peptidomimetic Cy-9B as an orange cartoon. The transparent yellow surface depicts the region defined by the funnel potential employed in the funnel-metadynamics calculations. **(B)** *Top:* Binding free energy surface (bFES) as a function of the distance between the centers of mass of the eIF4E binding pocket and Cy-9B, and the torsion angle describing the peptide’s orientation relative to the pocket. *Bottom:* One-dimensional projection of the bFES along the distance coordinate. The binding pose corresponding to the global free-energy minimum is shown in panel **C**. **(C)** Representative binding mode of Cy-9B at the global minimum, viewed from the top and the side. All Cy-9B side chains are shown in licorice representation, while only the key interacting residues of eIF4E are displayed. Black dashed lines indicate major interactions. **(D)** Superposition of the Cy-9B-eIF4E complex with the CYFIP1 crystal structure (chain A, PDB ID: 3P8C). Cy-9B aligns remarkably well (<1 Å) with the corresponding region of CYFIP1 used as its design template, demonstrating how effectively the peptidomimetic reproduces the native CYFIP1-eIF4E intraction.

To describe the binding process between Cy-9B and eIF4E, two CVs were used. The first is the distance r defined by the center of mass (COM) of residues selected at the binding pocket of eIF4E and the COM of selected residues in Cy-9B. The second CV is the angle θ that describes the rotation of one system over the other. This two-dimensional CV setting (r, θ) resembles the polar coordinates system found in other applications in many branches of science and mathematics (Fig. S3) ^54,55^. The bFES calculated as a function of the two CVs shows the lowest energy minimum at CV values 1.15 nm and 1.45 rad (Fig. 2B). Here, the binding mode of Cy-9B to eIF4E is characterized by the parallel orientation of Cy-9B’s α-helix with respect to the two α-helices forming the eIF4E binding pocket (Fig. 2C). In this pose, three salt bridge interactions are engaged by Cy-9B and eIF4E. In detail, Cy-9B’s Arg6, Arg8 and Glu10 interact with eIF4E’s Glu133, Glu71 and Arg129, respectively. An additional H-bond between Cy-9B’s Lys12 and eIF4E’s Asn78 further stabilizes the Cy-9B/eIF4E complex. The binding free energy ΔG calculated as the free-energy difference between the lowest energy binding mode and the unbound state is -11.6 kcal/mol, which corresponds to a dissociation constant (Kd) of ∼ 3 nM (Fig. 2B). The structural and energetic stability of the Cy-9B/eIF4E binding complex was further assessed by 200 ns of plain MD calculations (see Supplementary Information). It is worth noting that the Cy-9B binding mode is different from that reported for other 4E-BPs ^18,22,56–60,68,69^. In these structures, the α-helical region of the 4E-BPs assumes a perpendicular orientation relative to the α-helices forming the eIF4E binding pocket, interacting with different residues with respect to Cy-9B.

Prompted by our results and given the lack of an experimental structure of the CYFIP1-eIF4E complex, we docked CYFIP1 to eIF4E by aligning the CYFIP1’s chain A, obtained from the experimental structure with PDB ID 3P8C, to Cy-9B in the FM binary complex with eIF4E (Fig. 2D). Remarkably, residues 721-734 of CYFIP1 well overlap with residues 1-14 of Cy-9B, with a low RMSD value of 0.97Å calculated for the backbone atoms, showing a similar position for the residues involved in the binding interaction with eIF4E such as Arg726, Arg728, Glu730 and Lys732 (Arg6, Arg8, Glu10 and Lys12 in Cy-9B, respectively).

### Synthesis of CYFIP1-derived peptides and *in vitro* characterization

We synthesized Cy-9B along with its linear analogue Cylin and a scrambled peptide as control (Table S1). Circular dichroism (CD) spectroscopy corroborated the computational predictions (Fig. 1 and 3A), demonstrating that Cy-9B adopts a predominantly α-helical conformation in solution, an architecture required for productive engagement with eIF4E and essential for its pharmacological activity^21^. In contrast, the linear peptide (Cylin) was largely unstructured, exhibiting the spectral characteristics of a random coil (Fig. 3A). The stability of the Cy-9B peptide was then evaluated in human serum by comparison with its unmodified linear precursor. Peptide-based therapeutics often suffer from rapid proteolytic degradation, resulting in short circulatory half-lives that limit their clinical applicability (*i.e.*, minutes) ^70^, mainly due to enzymatic degradation. Notably, the unmodified linear peptide (depicted in blue) was completely degraded within just 1 hour of incubation with human serum proteases (Fig. 3B). In contrast, the stapled Cy-9B peptide (depicted in green) exhibited remarkable stability, with approximately 70% remaining even after 32 hours of incubation (Fig. 3B). Next step in the characterization of Cy-9B activity was to assess its binding to the eIF4E protein *in vitro*. The full-length human His-tagged eIF4E recombinant protein was produced in *E. coli* BL21 (DE3) pLysS cells, and its ability to bind the m^7^GTP cap was confirmed to verify proper folding and functional integrity (Fig. S4). Subsequently, the interaction between eIF4E and a biotinylated form of Cy-9B (Table S1) was analyzed by Bio-Layer Interferometry (BLI). The assay yielded a Kd of 57 nM for Cy-9B (Fig. 3C), corresponding to -9.94 kcal/mol, in line with the value predicted by FM (-11.6). Compared to the stapled peptide, the linear peptide exhibited a weaker affinity, with a Kd of 113 nM (Fig. 3D). A scrambled control peptide was designed based on the same primary sequence, with key residues critical for eIF4E binding purposely mutated, to disrupt specific interaction while maintaining overall amino acid composition. This peptide showed no detectable binding, underscoring the high specificity of Cy-9B toward eIF4E (Fig. S5A-B). Specific binding between Cy-9B and eIF4E was further corroborated by pulldown experiments (Fig. S5C-D). Prompted by these findings, we next examined the functional and cellular activity of Cy-9B. As an initial step, we assessed its ability to penetrate the cell membrane using the human lung cancer cell line A549 as a model system. Cy-9B, Cylin and the scrambled peptide (Fig. 3E, and Table S1) were labeled with fluorescein-maleimide and analyzed for cellular uptake and intracellular localization by flow cytometry and confocal microscopy (Fig. 3F-J). After 2h of incubation, cellular uptake of fluorescein-labeled Cy-9B was dose-dependent: ∼50% of A549 cells were fluorescent at 10 µM, 93% at 20 µM, and >98% at 40 µM, relative to untreated controls (Fig. 3G). Encouraged by these results, we selected this concentration for a time-course analysis of A549 cells treated with fluorescein-labeled Cy-9B and scrambled control peptides over 0–120 minutes. Flow cytometry revealed a gradual increase in intracellular fluorescence for both peptides. Notably, while the scrambled peptide also showed uptake, its fluorescence plateaued at just over 50%, compared with approximately 90% for Cy-9B (Fig. 3H). Moreover, the Cylin peptide displayed uptake patterns comparable to the control, with significantly reduced internalization relative to the stapled Cy-9B (Fig. 3I). These results are consistent with the well-established notation that peptidomimetics adopting an α-helical conformation typically exhibit enhanced cellular membrane penetration compared with unstructured linear peptides ^71,72^. To further evaluate uptake, cells were treated with 20 µM fluorescein-labeled Cy-9B for 6 hours. This prolonged exposure ensured maximal internalization and allowed assessment of subcellular localization beyond the plasma membrane. Confocal imaging (Fig. 3J) revealed predominantly cytoplasmic distribution, confirming effective intracellular entry.

**Fig. 3.**
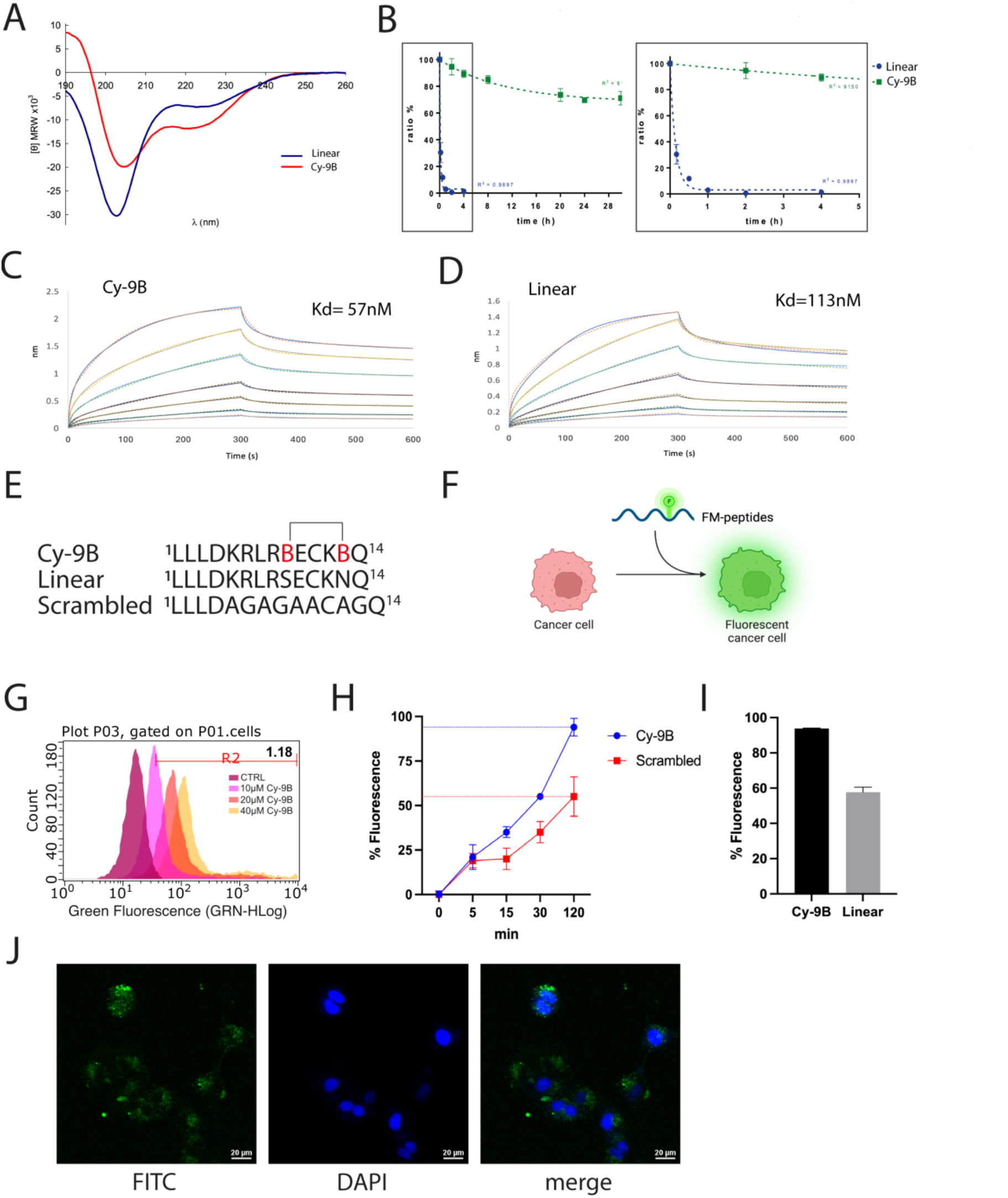
*In vitro* characterization of CYFIP1-derived peptides. **(A)** Circular Dichroism (CD) spectra of the linear peptide Cylin (blue line) and Cy-9B (red line). The Cy-9B exhibits the characteristic signature of an α-helix, whereas the linear peptide displays a random coil profile. **(B)** Human serum stability assay. *Left*: the degradation percentage of the peptides over 30-hour time course. *Right*: Expanded view of the first 5 hours, highlighting the fast degradation of the linear peptide in blue compared with Cy-9B in green. Data represent mean ± SD of three replicates. **(C-D)** Bio-Layer Interferometry binding kinetics assay. Binding affinity of Cy-9B **(C)** and Cylin **(D)** at different concentrations of eIF4E. Kds were calculated by steady-state fitting of maximum BLI responses. Representative sensorgrams and global fits are shown. Experiments were performed in triplicate. **(E)** Sequence of Cy-9B, lin-Cy-9B and the scrambled peptide, each containing a cysteine residue for fluorescein–maleimide labeling. (**F**) Schematic overview of the cellular uptake workflow. **(G)** Flow cytometry analysis performed on human lung cancer cells after 2h of incubation with fluorescein-labeled Cy-9B. Peaks correspond to untreated cells (magenta) and Cy-9B at 10 µM (pink), 20 µM (orange), and 40 µM (yellow). **(H)** Time-course analysis (0-2 h) of Cy-9B (blue line) and scrambled peptide (red line) uptake in A549 cells by flow cytometry. **(I)** Comparison of cellular fluorescence after 2 h of incubation with Cy-9B versus Cy-lin. **(J)** Confocal microscopy images of A549 cells treated with fluorescein-labeled Cy-9B. Cy-9B (green) localizes predominantly to the cytoplasm. Nuclei are stained with DAPI (blue). The merged image confirms effective intracellular internalization of Cy-9B.

### Cy-9B disrupts eIF4F assembly and inhibits cap-dependent translation in a lung cancer cell model

eIF4E is overexpressed in a wide variety of human cancers (Fig. S6A). Although CYFIP1 can also be upregulated in certain tumor types (Fig. S6B), its expression is comparatively low in lung cancer (Fig. S6C), suggesting weakened endogenous repression of eIF4E and heightened reliance on cap-dependent translation. Consistent with this, CYFIP2, sharing 88% amino acid identity with CYFIP1, was recently shown to have prognostic value in lung cancer (Fig. S6D) ^73^. Together, these data underscore the relevance of the eIF4E-CYFIP1 axis in this malignancy and support eIF4E as a therapeutic target.

To evaluate whether Cy-9B modulates eIF4F assembly, we performed a m^7^GTP Sepharose pull-down assay in A549 lung cancer cells derived lysates (Fig. 4A), a standard method for investigating the composition of the eIF4F complex ^18,24,74–76^. In untreated cell lysates, eIF4G co-purified with eIF4E as expected. Cy-9B treatment for 2 h results in a dose-dependent decrease in eIF4G bound to eIF4E (Fig. 4B), with ∼50% reduction at 50 µM Cy-9B (Fig. 4C), consistent with Cy-9B competing with eIF4G for the canonical eIF4E-binding interface. To determine the functional consequences of this disruption, we transfected A549 cells with a plasmid encoding a bicistronic reporter in which Renilla luciferase is translated via a cap-dependent mechanism, whereas Firefly luciferase is translated via a cap-independent internal ribosome entry site (IRES). The day after transfection, cells were treated with increasing concentrations of Cy-9B and incubated for 2 hours, displaying selective inhibition of Renilla activity, with ∼50% suppression at 25-50 µM (Fig. 4D). Firefly luciferase, representing cap-independent translation, was unaffected or modestly increased, in agreement with previous observations ^37^. These findings confirm that Cy-9B specifically impairs the eIF4E-eIF4G dependent initiation pathway.

**Fig. 4.**
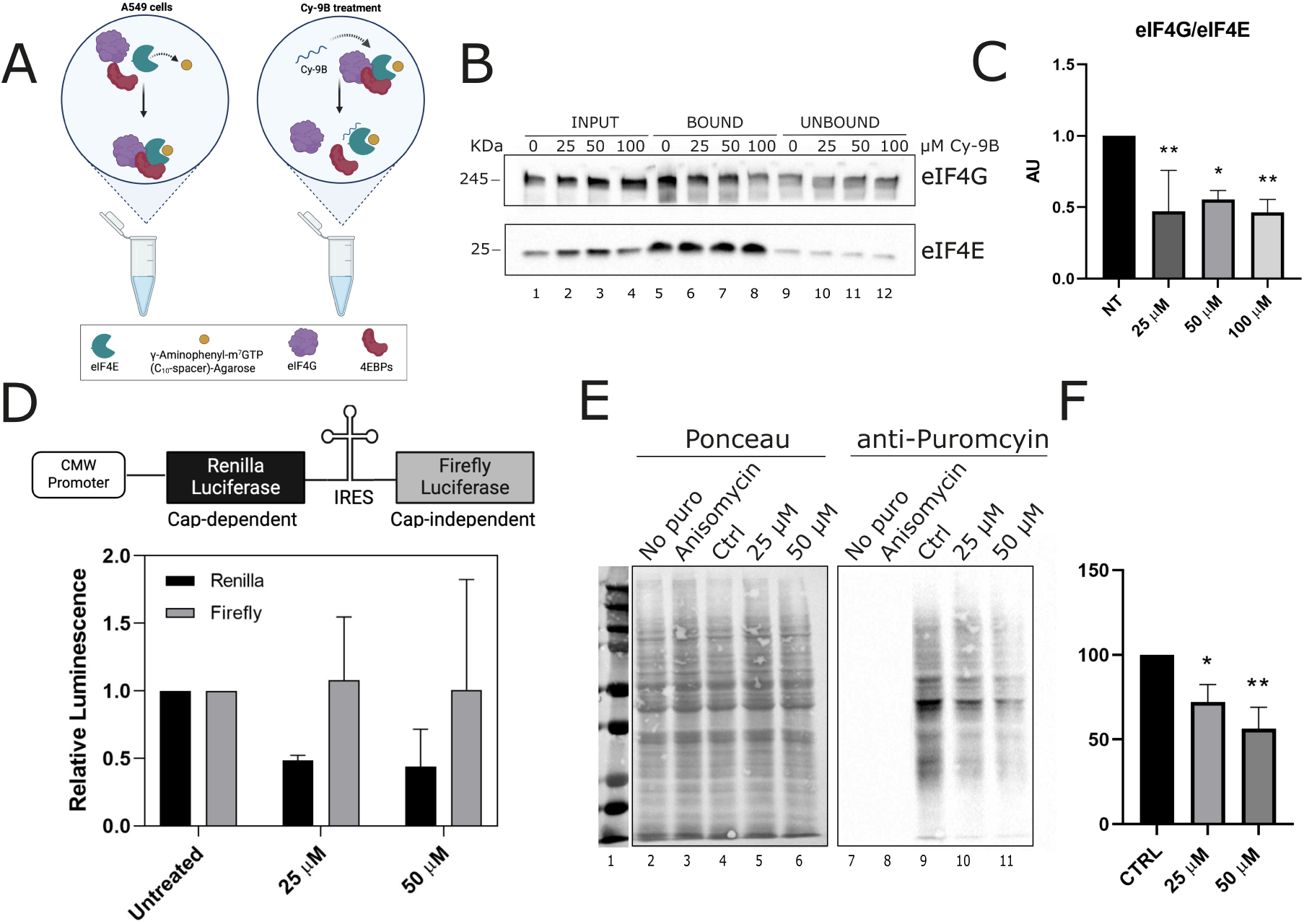
Cy-9B inhibits cap-dependent translation in A549 lung cancer cells. **(A)** Schematic overview of the m^7^GTP pull-down assay. **(B)** Representative Western blot and **(C)** corresponding quantification of m^7^GTP pull-downs performed on A549 cell extracts treated with Cy-9B at the indicated concentrations. Lanes 1-4: input (1/20); lanes 5-8: bound fraction; lanes 9–12: unbound fractions. m^7^GTP chromatography was conducted in presence of 0 µM (vehicle, NT; lanes 1, 5, 9), 25 µM (lanes 2, 6, 10), 50 µM (lanes 3, 7, 11), and 100 µM (lanes 4, 8, 12) Cy-9B. Data represent mean ± SEM, normalized to vehicle controls, and analyzed using two-way ANOVA. **(D)** Schematic of the pcDNA3-Rluc-Pol IRES-FLuc bicistronic construct (top). Renilla (cap-dependent) and Firefly (IRES-dependent) luciferase activities in A549 cells were measured after 2 h of Cy-9B treatment (bottom) and expressed relative to untreated controls (bottom). **(E)** Western blot analysis of global protein synthesis using the SUnSET puromycin-incorporation after 24h treatment with Cy-9B at different concentrations. **(F)** Quantification of puromycin incorporation normalized to Ponceau staining. Data are presented as fold change relative to untreated cells, represented as mean ± SEM from triplicate. Statistical significance was determined by one-way ANOVA.

Given this targeted effect, we next assessed global translation using the SUnSET puromycin-incorporation assay after 24 h of peptide treatment. Cy-9B reduced overall protein synthesis to ∼30% of control levels at 50 µM (Fig. 4E-F), whereas the scrambled peptide had no appreciable effect (Fig. S7A). These results establish that Cy-9B effectively inhibits eIF4F complex formation and suppresses cap-dependent protein synthesis in lung cancer cells.

### Cy-9B suppresses tumor cell growth, adhesion, migration, and invasion

We next measured the impact of Cy-9B on A549 lung cancer cell growth, using the scrambled peptide as control. Cell viability and proliferation were assessed by MTT (3-(4,5-dimethylthiazol-2-yl)-2,5-diphenyltetrazolium bromide) assay after treatment with 25, 50, and 100 µM of Cy-9B or scrambled peptide for 24, 48, and 72 hours. Peptides were added once at the start of the experiment and maintained for the entire duration without medium replacement. Cy-9B reduced cell viability in a dose-dependent manner, with an IC50 of 64 µM at 24 h, 67 µM at 48 h, and 87 µM at 72 h (Fig. 5A and Fig. S7C). In contrast, the scrambled peptide showed no effect (Fig. 5B). These data indicate that Cy-9B impairs proliferation of A549 cells, with maximal inhibitory potency after 24h, as reflected by the lowest IC50 value, and does so without inducing cell death (Fig. S7B). Together with our mechanistic analyses, these results support a model in which Cy-9B suppresses cell growth by targeting cap-dependent translation.

**Fig. 5.**
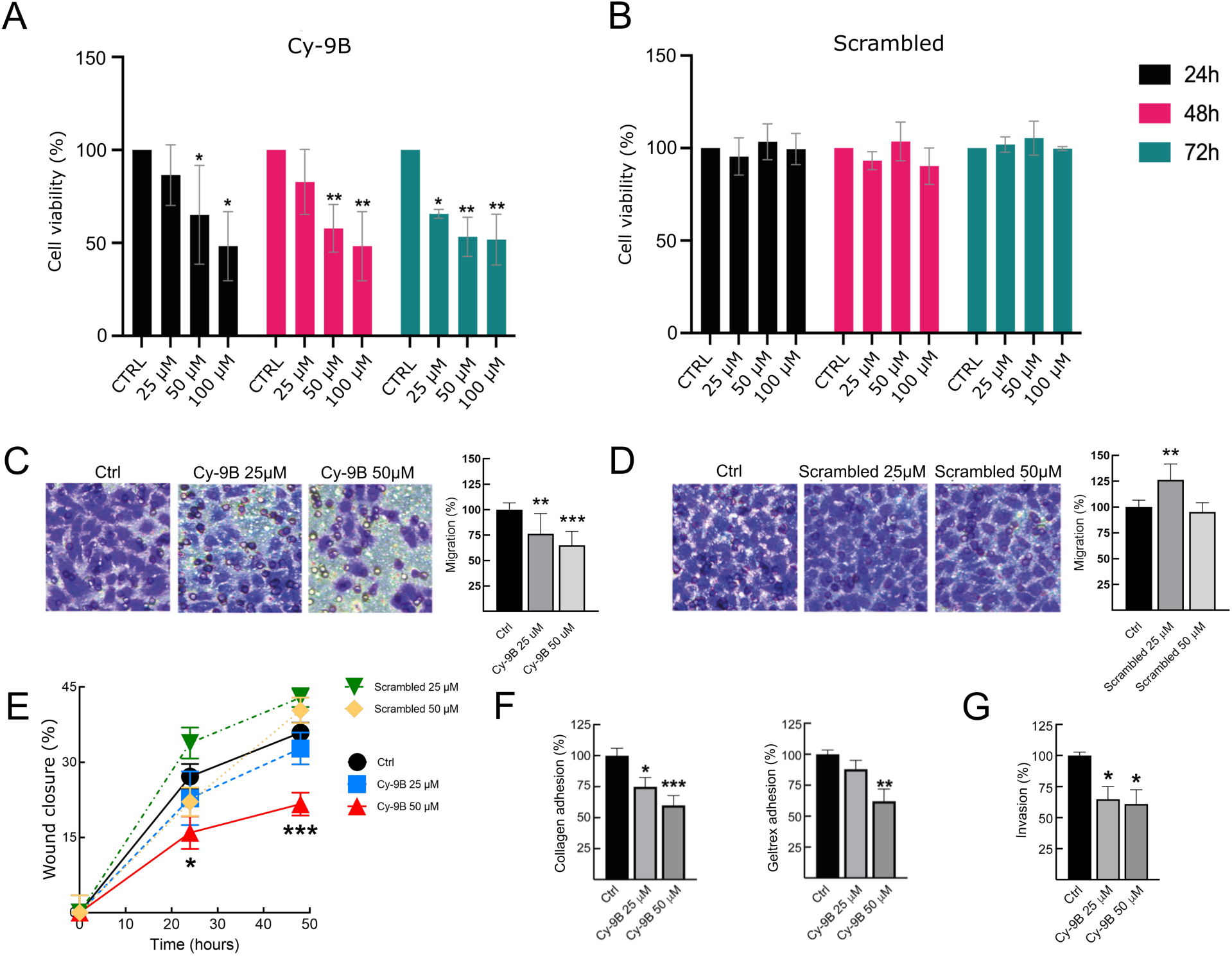
Cy-9B peptide impairs lung cancer cell proliferation, migration, and invasion. **(A-B)** Growth inhibition of A549 cells assessed by MTT assay. Cell viability of human lung cancer cells after 24 h (in black), 48 h (in magenta), and 72 h (in green) of treatment with increasing concentrations of Cy-9B **(A)** or scrambled peptide **(B)** as a control. Experiments are triplicate and data are mean ± SEM. Statistical analysis was carried out using one-way ANOVA with Tukey’s multiple comparison post hoc test versus to untreated control. **(C-D)** Cy-9B inhibits A549 cell migration in Boyden chamber assay. A549 cells were pre-treated for 24 h with 25 or 50 µM Cy-9B **(C)** or scrambled peptides **(D)**, then, seeded into the upper chamber with the same peptide concentrations present in the assay. After 24 hours, migrated cells were stained with Crystal violet; dye was solubilized and absorbance measured at 590 nm. Migration was quantified as a percentage relative to control, as shown in the accompanying graph. Data are mean ± SEM (One-Way ANOVA followed by Tukey’s multiple comparison test). **(E)** Wound-healing assay showing reduced A549 cell migration after treatment with 25 or 50 µM Cy-9B or scrambled peptide for up to 48 h. Migration was quantified by measuring the distance between the wound edges in more than 3 randomly selected fields per condition. Data are mean ± SEM (One-Way ANOVA followed by Dunnett’s multiple comparison test). **(F)** Effect of Cy-9B on A549 cell adhesion to extracellular matrix components. A549 cells (4×10^5^/ml) were seeded to individual coated wells with 200 µl of collagen (7.5 µg/ml) (left panel), or geltrex (0.2 mg/ml) (right panel), in absence or in presence of 25 and 50 µM Cy-9B and incubated for 30 minutes at 37°C in 5% CO_2_. Adhesion was quantified and expressed as percentage relative to untreated control cells. (mean ± SEM; n ≥ 3, one-way ANOVA followed by Tukey’s multiple comparison test. **(G)** Cy-9B reduces the invasive capacity of A549 cells. Cells were pre-treated for 24 h with 25 and 50 µM Cy-9B and then seeded in the upper part of a Boyder chamber coated with collagen I, in the presence and in the absence of Cy-9B 25 and 50 µM. After 24 h, invaded cells were stained with crystal violet, solubilized with ethanol, and absorbance read at 590 nm. Data are mean ± SEM (n ≥ 3; One-Way ANOVA followed by Tukey’s multiple comparison test).

Given the established role of eIF4E in promoting malignant phenotypes, including migration, invasion, cell–matrix adhesion, and epithelial-to-mesenchymal transition ^77,78^, we next investigated whether Cy-9B modulates these cancer-associated behaviors. We first tested the effect of Cy-9B and the scrambled peptide on A549 lung cancer cell migration in Boyden chamber assays (Fig. 5C-D). The results indicated that Cy-9B significantly reduced cell migration at 25 µM and showed an even stronger inhibitory effect at 50 µM (Fig. 5C-D). Consistently, wound-healing assays revealed that Cy-9B markedly delayed scratch closure after 24 h, with further suppression at 48 h, reaching ∼20% inhibition of migration (Fig. 5E and S7D-E). To evaluate effects on cell-matrix adhesion, A549 cells were pre-treated for 24 h with 25 or 50 µM Cy-9B and plated onto either collagen or geltrex. The latter is a basement membrane matrix which includes laminin, collagen IV, entactin, and heparin sulfate proteoglycans, mimicking the extracellular matrix efficiently. The results indicated that Cy-9B significantly impaired adhesion to both substrates. Specifically, at 25 µM Cy-9B, adhesion to collagen decreased by ∼25%, and at 50 µM by ∼40%. Similar inhibition was observed on geltrex (Fig. 5F and S7F).

Finally, using a collagen I-based invasion assay, we found that Cy-9B reduced the invasive capacity of A549 cells by ∼35%. This effect was statistically significant already at 25µM Cy-9B (Fig. 5G and S7G).

### Cy-9B reduces global protein synthesis in an *in vitro* FXS model and rescues social behavior in *Cyfip*^85.1^*/+* flies

As noted above, dysregulated protein synthesis is a convergent feature of diverse pathologies, including cancer and FXS ^79^. In tumors, this imbalance is often driven by elevated eIF4E activity, whereas in FXS it arises from disruption of the FMRP-containing translational repression machinery. Whether driven by eIF4E overexpression, as in many cancers, or by disruptions of the FMRP-containing translational repression complex in FXS, the pathological outcome converges on a common imbalance of the eIF4E-CYFIP1-FMRP axis, resulting in aberrant activation of the cap-dependent translation (Fig. 6A). Despite extensive research, no therapy has yet been approved for FXS. Several candidates are currently in clinical evaluation, including the PDE4D inhibitor zatolmilast, the BK channel activator SPG601, and the IGF-1-derived peptide analog trofinetide^80^, yet, to date, no peptidomimetic has progressed to clinical trials^12^.

**Fig. 6.**
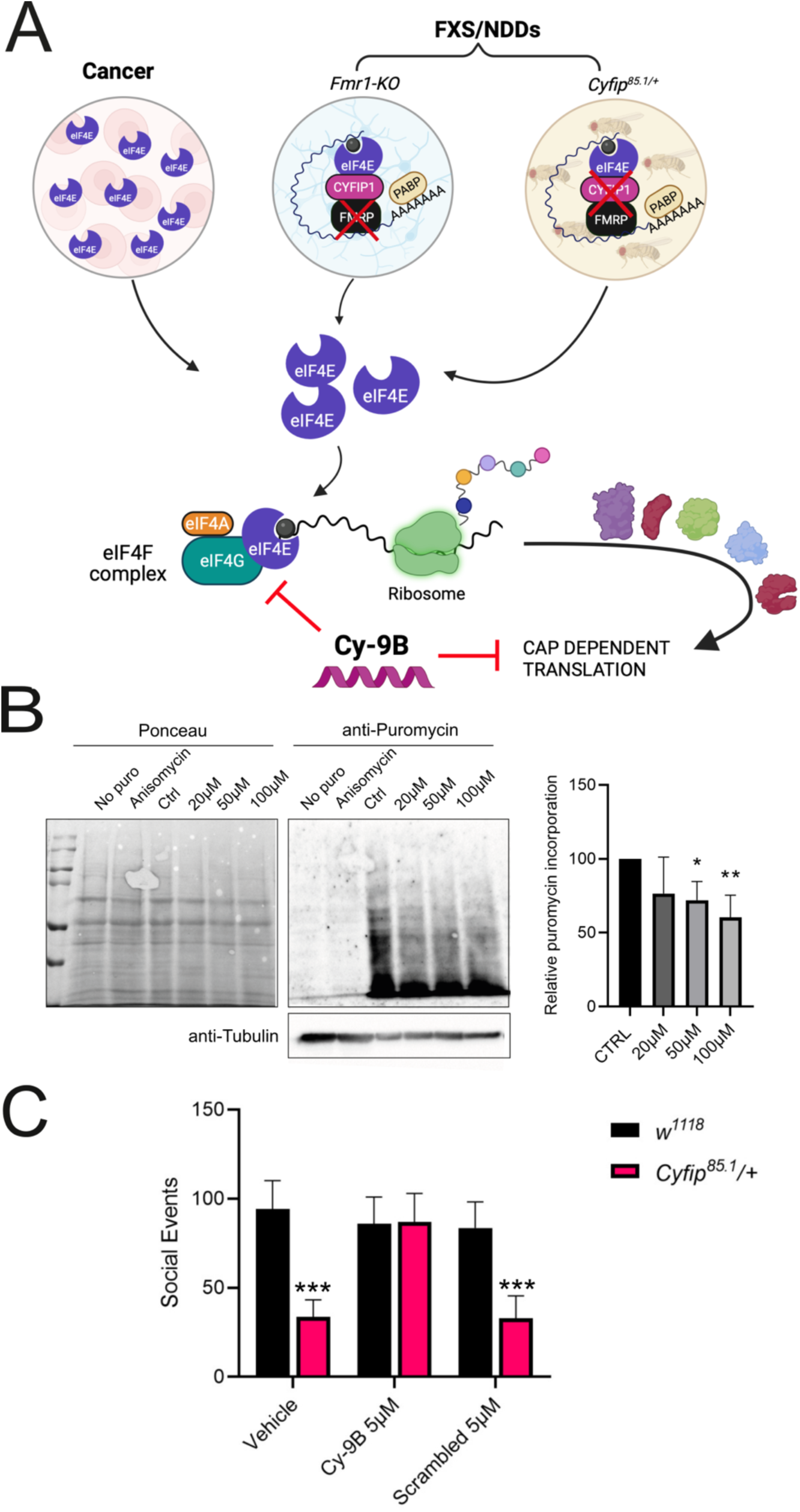
Dysregulation of cap-dependent translation in cancer and neurodevelopmental disorders (NDDs). (**A**) Conceptual schematic summarizing disease-specific perturbations that converge on enhanced eIF4E-driven translation. *Left*: in cancer, eIF4E overexpression promotes elevated cap-dependent protein synthesis. *Middle*: in FXS, loss of FMRP in hippocampal Fmr1-KO neurons disrupts the FMRP-containing translational repression complex, favoring eIF4E complex formation and increased translation. *Right*: *Cyfip^85.1^/+* Drosophila, a CYFIP haploinsufficiency model relevant to CYFIP1-associated NDDs (*e.g.*, ASD, schizophrenia, and intellectual disability), exhibits impairment of the CYFIP1-FMRP-eIF4E regulatory axis, similarly shifting translation toward hyperactivation. *Bottom*: schematic of cap-dependent translation initiation. eIF4E is normally released from inhibitory 4E-binding proteins (4E-BPs), including CYFIP1, in a phosphorylation-dependent manner. After release, eIF4E binds to the mRNA 5’-cap, and subsequently forms the eIF4F complex with helicase eIF4A and scaffold protein eIF4G. This complex promotes ribosome recruitment and initiation of cap-dependent translation. The proposed site of action of Cy-9B, disruption of eIF4E–eIF4G engagement, is indicated. **(B)** Global protein synthesis in *Fmr1*-KO neurons was measured by the SUnSET assay. Representative immunoblot showing Ponceau staining, puromycin incorporation, and tubulin levels after 24h Cy-9B treatment at 0, 20, 50, and 100 µM. Relative puromycin incorporation was quantified using Image Lab, normalized to Ponceau staining, and expressed as fold change relative to untreated controls. The experiment was reproduced in five replicates, n=5, and data are mean ± SEM. Statistical significance was determined by one-way ANOVA. **(C)** Social behavior (competition for food) of *Cyfip^85.1^*/+ flies upon treatment with 5 µM of Cy-9B or scrambled peptides for 12 hours. Aggressive events were quantified during a 2-min observation period in the presence of a food droplet, following 90 min of starvation and 2 min of re-exposure to food (habituation). n = 10 groups (8 flies per group) per genotype, tested across 3 independent experiments. *** p < 0.0001 (two-way ANOVA with Tukey’s multiple-comparisons test).

Given this therapeutic gap, and the convergence of distinct pathological mechanisms into shared CYFIP1-centered translational hub, we evaluated whether Cy-9B could modulate translation and its functional outcome across complementary models of FXS-associated dysregulation. As a first step, we assessed the impact of Cy-9B on translation in an in vitro neuronal model of FXS. Potential Cy-9B toxicity was evaluated in wild-type hippocampal neurons through MTT assay, since our earlier characterizations were performed in non-neuronal cells. Hippocampal neurons exposed to Cy-9B concentrations up to 100 µM for 24h, and the assay revealed no detectable toxicity for neuronal cells (Fig. S8A).

We next tested the effect of Cy-9B in neurons derived from *Fmr1-Knockout* mice (*Fmr1*-KO), a well-established FXS model characterized by elevated protein synthesis ^81^. Neurons were treated for 24 h with Cy-9B or scrambled peptide as control, and global protein syn-thesis levels were quantified using the SUnSET assay (Fig. 6B and S8B). Total protein content was assessed by Ponceau staining, and nascent protein synthesis was measured by puromycin incorporation detected with an anti-puromycin antibody. Anisomycin (40 µM, 15 min pre-treatment) was used as a positive control for translation inhibition. Cy-9B treatment (50 and 100 µM) reduced protein synthesis by more than 25% (Fig. 6B), whereas scrambled peptide had no effect (Fig. S8B).

Finally, we asked whether restoring control of the eIF4E-CYFIP1 axis by Cy-9B yields organism-level functional benefit. To this end, we administered Cy-9B *in vivo* to *Cyfip*^85.1^*/+ Drosophila melanogaster*, and quantified social behavior. Consistent with prior reports, CYFIP1 haploinsufficient flies exhibit ASD-relevant phenotypes, including alterations in social and repetitive behaviors^32^, and associated comorbidities such as sleep abnormalities^34^. In the social dominance paradigm^32^, Cyfip heterozygous flies display significantly reduced aggression following starvation and subsequent limited food re-exposure, indicative of im-paired social dominance relative to controls (P<0.0001) (Fig. 6C). Notably, Cy-9B administration rescued this deficit, restoring aggression to control-like levels (P=0.8564). By contrast, the scrambled peptide failed to ameliorate the phenotype (P<0.0001), supporting the specificity of Cy-9B–mediated rescue.

## Discussion

Our findings establish Cy-9B as a rationally engineered, CYFIP1-inspired peptidomimetic with broad therapeutic promise. Designed to target eIF4E, Cy-9B was characterized using complementary computational and experimental approaches, including funnel metadynamics, which revealed a non-canonical binding mode that differs from previously described 4E-BP-derived ligands. This distinct recognition profile provides a new structural template for achieving selectivity within the eIF4E interactome^82^. In our previous work^21^, we proposed an initial binding mode for CYFIP1 peptide to eIF4E. Leveraging recent methodological advances, the present study refines this model, revealing additional contact residues engaged by Cy-9B with eIF4E leading to a specific interaction interface not exploited by canonical 4E-BP motifs ^82^.

Because CYFIP1 engages eIF4E through an α-helical motif, we stabilized this conformation through a triazole-bridged staple to minimize the entropic cost of folding upon binding while improving proteolytic stability and cellular uptake. Consistent with this rationale design, Cy-9B exhibited increased structural stability relative to its linear precursor, higher affinity for eIF4E, and potent disruption of the eIF4E–eIF4G complex, an essential gatekeeper of cap-dependent translation initiation. Functionally, Cy-9B reduced global protein synthesis and impaired proliferation, migration, and invasion in lung cancer cells, while also attenuating elevated translation in FXS hippocampal neurons. Across systems, 50 µM emerged as the most consistently effective concentration. Importantly, Cy-9B operates as a partial translation inhibitor rather than a global suppressor. Such calibrated attenuation is advantageous given the essential role of protein synthesis in viability, and mirrors the therapeutic logic of other eIF4E-targeting strategies that seek modulation rather than ablation of translation^13^.

In oncology, this mechanism offers a targeted alternative to broadly cytotoxic approaches. In FXS, it introduces a conceptually distinct therapeutic strategy: direct restoration of translational homeostasis by re-engaging a core molecular brake on eIF4E activity. To date, peptides and peptidomimetics have not been explored as a means to restore FMRP-dependent translational control in FXS. By acting as a functional surrogate for the CYFIP1-FMRP-eIF4E regulatory module, Cy-9B has the potential to address a proximal pathogenic mechanism rather than downstream symptoms^72^. This is particularly relevant given that current FXS treatments remain symptomatic and lack disease-modifying efficacy^83,84^. Emerging clinical approaches, such as mavoglurant (mGluR5 antagonists), arbaclofen (GABAergic modulators), experimental gene therapies, and anti-sense oligonucleotides (ASOs) ^85^, have faced limitations in consistency and clinical applicability. Similarly, other candidates, such as zatolmilast (BPN14770, a phosphodiesterase-4D inhibitor) ^86^, Zygel (ZYN002, a cannabidiol transdermal gel) ^87^, both in Phase 3 clinical trial, and CTH120 (<u>a “neuroplasticity modulator” developed using artificial intelligence and currently in Phase 1 clinical trial</u>), offer symptomatic benefit but do not directly target the dysregulated translation machinery. Cy-9B’s translational relevance is further reinforced by organism-level rescue in an *in vivo* genetic model. In *Cyfip*^85.1^*/+ Drosophila melanogaster*, which display robust deficits in social behavior, Cy-9B restored social interaction phenotypes to wild-type-like levels, whereas a scrambled control peptide was ineffective.

The lower effective concentration *in vivo* in Drosophila (5 µM, behavioural readout) compared with neuronal culture (50–100 µM; molecular readout) is consistent with the perturbation acting across distributed circuits, where shifting activity in a small set of nodes can recruit larger networks and produce a discrete change in behaviour ^32,34,88,89^. Together, these results demonstrate Cy-9B as a functional surrogate of the CYFIP1-FMRP-eIF4E translational brake, linking molecular target engagement to translational control and, ultimately, to cross-species behavioral outcomes, thereby strengthening the case for eIF4E-centered modulation as a disease-modifying strategy in FXS-related NDDs.

Looking ahead, Cy-9B provides both a first-in-class lead compound and a mechanistic framework for structure-based optimization of selective eIF4E modulators. In the context of FXS, such agents may be most impactful if deployed during defined developmental windows of synaptic circuit formation, potentially maximizing durable benefit while minimizing long-term exposure. More broadly, our work highlights a shared vulnerability, hyperactivated cap-dependent translation, across cancer and NDDs, and positions selective, physiologically compatible tuning of eIF4E activity as a generalizable therapeutic avenue.

## Methods

### Peptide synthesis

Peptides were synthesized by microwave-assisted Fmoc-SPPS on a Biotage ALSTRA Initiator+ peptide synthesizer, operating in a 0.12 mmol scale on a Rink amide resin (C-terminus amidated peptides). Resins were swelled prior to use with a NMP/DCM mixture (1:3) for 1 hour. Activation and coupling of Fmoc-protected amino acids were performed using Oxyma 0.5M / DIC 0.5M (1:1:1), with a 4-equivalent excess over the initial resin loading. Cysteine residues were included as S-Acm protected, and the Acm group was removed after click-based macrocyclization (see below). Coupling steps were performed for 10 minutes at 60°C. Deprotection steps were performed by treatment with a 20% piperidine solution in DMF at room temperature (1 × 5min). Following each coupling or deprotection step, peptidyl-resins were washed with DMF (4×). Peptides were cleaved from the resin using a TFA 90%, water 5%, thioanisole 2.5%, TIS 2.5% mixture (2 hours, RT), and then precipitated in cold diethyl ether. Crude peptides were collected by centrifugation and washed with further cold diethyl ether to remove scavengers. Linear precursors were high-performance liquid chromatography (HPLC) analyzed and purified as described below.

#### Intramolecular Cu(I) -catalyzed azido-alkyne 1,3-cycloaddition for peptide stapling

To the linear peptide precursor solutions (0.5 mg/ml in degassed water) CuSO_4_. 5 H_2_O (5 eq) and ascorbic acid (5 eq) were added to generate Cu(I) catalyst *in situ*. Stapling reactions were stirred at room temperature until complete conversion of the linear precursor into the desired heterodetic 1,2,3-triazolyl-containing peptide, as monitored by reversed-phase high-performance liquid chromatography (RP-HPLC). The resulting stapled peptide was purified by RP-HPLC as described below.

#### Cys(Acm) deprotection

Following cyclization, the peptide was incubated in the presence of PdCl2 (5 eq., RT) until complete removal of the Acm group. Then, the reaction was quenched with diethyldithiocarbamate (DTC), and the deprotected compound was recovered by RP-HPLC purification.

### RP-HPLC analysis and purification

Analytical and semi-preparative reversed-phase high-performance liquid chromatography (RP-HPLC) were carried out on a Shimadzu Prominence HPLC system equipped with a multichannel detector. A Phenomenex Jupiter 5µ C18 90Å column (150 × 4.6 mm) was used for analytical runs and a Phenomenex Jupiter 10µ C18 90Å (250 × 21.2 mm) for peptide purification. Data were recorded and processed with LabSolutions software. 5-100 % linear gradient eluent B at a flow rate of 0.5 mL/min was used for analytical purposes (20 min run). eluent A = H_2_O/ 3 % CH_3_CN / 0.07 % TFA, eluent B = 70 % CH_3_CN/ 30 % H_2_O/ 0.07 % TFA. UV detection was recorded in the 220-340 nm range. Peptide purification was performed by preparative RP-HPLC at a flow rate of 14 mL/min using a 100% A→30% B linear gradient over 40 min. Pure fractions (>95%) were combined and lyophilized. LC-MS analysis was then performed on pure fractions.

### Circular Dichroism (CD) spectroscopy

CD spectra were collected on a Jasco J-815 spectropolarimeter equipped with a Peltier temperature control system. Peptide concentration was 30 µM in phosphate buffer, 20 mM, NaF 100 mM, pH 6.5. Measurements were acquired in a 1 mm quartz cuvette, at 280 K, using an average of four scans between 190 and 260 nm, with a scanning speed of 20 nm/min, 0.5 s of data integration time, and a resolution of 0.1 nm.

### Peptide stability assay in human serum

Serum stability assays were performed according to the protocol reported by Zhang et al. ^90^. Briefly, human serum (Sigma-Aldrich, H4522) was thawed and centrifuged (14,000 rpm at 4 °C for 10 min) before incubating at 37 °C for 15 min to activate the proteins. Peptide solutions were prepared at 300 µM in PBS and added to human serum in a 1:4 (v/v) ratio, and the mixed solutions (final concentration 75 μM) were incubated at 37 °C. At predefined timepoints, 35 µL aliquots of each sample were collected, mixed with 15 µL of 10% trichloroacetic solution, and left for 10 min at 4 °C. An additional 35 µL of water was then added, and the samples were centrifuged at 14,000 rpm for 10 min at 4 °C. The supernatants were injected onto an analytical C18 column (Shimadzu Shimpack GWS C18 column, 5 micron, 4.6 mm i.d. × 150 mm), and the resulting HPLC data were processed using LabSolutions software. The residual percentage of peptide was quantified by integrating the peak area from the HPLC chromatograms at 214 nm. All peptides were tested in triplicate. The stability data of peptides were calculated with GraphPad Prism using nonlinear regression, one-phase decay.

### Bio-Layer Interferometry assay

Biotinylated peptides were immobilized using a 2.5 ng/µl solution in Sartorius Kinetics Buffer to Octet® Streptavidin (SA) Biosensor.

A dilution series of the protein dissolved in kinetics buffer from 1.8 µM to 0.025 µM was used to determine the protein-peptide affinity. The experiments were run using an Octet® R8. The data were plotted and fitted using Octet® analysis studio (v.12.2.2.26).

### Cloning, expression, and purification of eIF4E

The gene coding for human eIF4E (Uniprot: P06730) was cloned from the pHA-eIF4E (Addgene #17343) into the pMCSG7 expression vector using Ligation Independent Cloning (LIC) as described in Bruni et al. ^91^ and used to transform *Escherichia coli* BL21 pLysS cells. *heIF4E* gene was PCR amplified using primers suitable for LIC cloning (Forward: 5’-TACTTCCAATCCAATGCCATGGCGACTGTCGAACCG-3’; Reverse: 5’-TTATCCACTTCCAATGTTAAACAACAAACCTATTTTTAG-3’). The cloned eIF4E gene produces the full-length initiation factor plus ten histidine residues at the N-terminus. The cells expressing the 10×His-eIF4E were grown in LB medium at 37°C until OD= 0.4-0.6, and eIF4E induction was started with 0.5 mM IPTG, and the culture was placed in a shaker-incubator (New Brunswick Innova) overnight at 20°C. Cells were harvested by centrifugation, and the cell pellets were re-suspended in 50mM HEPES, pH 7.8, 100 mM KCl, 1mM EDTA, 2mM DTT, 10% Glycerol, Protease Inhibitor Cocktail (Merk Millipore), 0.5% Triton X-100. The sample was then sonicated for 30 min at 12,000 x g at 4°C. The induced protein was recovered as an inclusion body. The resulting pellet was resuspended in 50mM HEPES, pH 7.8, 1M Guanidine, 10% Glycerol, 2mM DTT, and centrifuged for 30 min at 30,000 × g at 4°C. This procedure was repeated twice. The remaining pellet was solubilized with 50mM HEPES pH 7.8, 6M Guanidine, 10% Glycerol, 2mM DTT, and centrifuged for 30 min at 43,000 × g at 4°C. The concentration of the denatured protein was adjusted to 0,1 mg/ml into refolding buffer consisting of 50 mM HEPES pH 7.8, 3M Guanidine, 10% Glycerol, 2mM DTT and the refolded protein was dialyzed into 50mM HEPES pH 7.8, 300mM KCl, 2mM DTT, 10% Glycerol and concentrated with Amicon Ultra 10k (Merk Millipore) up to 1mg/ml.

### Mammalian cell culture conditions

Human A549 lung adenocarcinoma cells (ATCC CCL-185) were cultured in DMEM-HG cell media supplemented with 10% Fetal Bovine Serum (FBS) (Corning), 2mM Glutamine, and penicillin/streptomycin. Cells were maintained in a 37°C humidified incubator with 5% CO_2_ atmosphere.

### M^7^GTP Pulldown Assay

Cells at ∼80% confluence were washed three times with ice-cold 1× PBS and scraped in lysis buffer containing 50 mM Tris-HCl (pH 7.4), 5 mM MgCl_2_, 100 mM KCl, 0.5% Triton X-100, and protease/phosphatase inhibitors. Cap-binding complexes were captured using m**^7^**GTP-agarose resin (Jena Bioscience). Equal amounts of protein lysates (500 µg) were incubated with 30 µL of m**^7^**GTP resin, pre-saturated with 0.1 mg/mL BSA, for 2 h at 4°C to enrich cap-binding proteins. After collecting the unbound fraction, beads were washed three times with 500 µL of wash buffer (50 mM Tris-HCl, pH 7.4, 5 mM MgCl_2_, 100 mM KCl, 0.5% Triton X-100), and bound proteins were eluted with 2× Laemmli sample buffer. Eluates were analyzed by SDS–PAGE followed by immunoblotting using eIF4E antibody (1:1000, #9742, CST) and eIF4G antibody (1:1000, #2617, CST). Input and unbound fractions (1/20) were included as controls.

### Streptavidin-Biotin Pulldown Assay

Biotinylated peptides were used as indicated in Table S1. Each peptide (30 µg) was incubated with purified human N-His-eIF4E (500 ng) for 2 h at 4°C in 300 µL buffer (300 mM NaCl, 50 mM Tris-HCl, pH 7.5, 0.1% Triton X-100). Streptavidin-conjugated magnetic beads (30 µL, Invitrogen) were pre-equilibrated with buffer containing 0.1% BSA for 15 min at room temperature and then incubated with the peptide–eIF4E mixture for 1 h at room temperature. Beads were washed three times with the same buffer, and bound proteins were eluted with 2× Laemmli sample buffer. Eluates were analyzed by SDS-PAGE and immunoblotted for eIF4E.

### Dual-luciferase assay

Twenty thousand A549 cells were plated per well in 96-well plates. Cells were transfected with 200 ng of the bicistronic reporter construct pcDNA3-RLucPoliresFLuc (Addgene 45642) using polyethyleneimine (PEI) (25K, Polysciences) at a 1:3 ratio. PEI was prepared as a stock solution at 1 mg/mL in deionized (DI) water for transfection. The cells were incubated for 24 hours and treated for 2 hours before the lysis, with different concentrations of Cy-9B peptide. Then, the cells were lysed in 1× passive lysis buffer (Promega), transferred to white-opaque 96-well plates and luciferase activity was measured with a dual luciferase reporter assay system (Promega) using an Infinite200 plate reader (Tecan Life Sciences). A non-transfected control cells were used to remove the luminescence background signal. Duplicate wells were set up for each assay, and assays were independently done three times.

### SUnSET Assay

Global de-novo protein translation was measured by using the SUnSET assay as previously described ^92^. In detail, A549 cells (100,000 cells/well) were seeded into 12-well plates. After 24h, cells were treated with 25 and 50 µM of peptides for another 24h. Then, cells were treated with 2 µM puromycin for 30 min, washed twice with PBS, and then lysed in Sample Buffer 3×. Pre-treatment with the translation inhibitor anisomycin (40 µM) for 15 min was used as a control to confirm the specificity of the assay. 15 µl of samples was subjected to SDS-PAGE and puromycin incorporation was analyzed by western blotting using an anti-puromycin antibody (1:10,000 clone 12D10, Merck Millipore). Ponceau red staining was used as a protein loading control. Experiments with hippocampal cells were carried out at DIV 13 in 12-well plates pre-coated with PLL, following the protocol previously described for A549 cells at a density of 2.5×10^5^ cells/well.

### Fluorescent labeling of peptides

Fluorescein-5-maleimide (FM) (Thermo Fisher Scientific) at 100 µM in 10% (v/v) dimethylformamide (DMF) in PBS, pH 7.4, was incubated with 400 µM of peptides dis^-^solved in PBS/EDTA 5mM for 2 hours at room temperature as in ^36,93,94^. The unreacted fluorescein was removed using Fluorescent Dye Removal Columns (Thermo Fisher Scientific). Labeled peptides were used at 400 µM stock concentration (corresponding to 100 µM final concentration) for cellular uptake experiments without further purification.

### Flow cytometry analysis

A549 cells were seeded into 24-well plates (25,000 cells/well) and incubated with peptides at varying concentrations and time points in 500 µL of medium. Cells were harvested by trypsinization, washed, and resuspended in PBS before subjecting them to flow cytometry analysis. Fluorescence peptide was measured with Guava® easyCyte™ Flow Cytometer (Cytek, Fremont, CA, USA) equipped with a 488nm laser. Peptide fluorescence intensity was detected with following gain settings: FSC:25.8 SSC: 12.9; G: 5.19; Y:1 R:1 R2: 1. A minimum of 5000 cells were acquired. For quantitative analysis, a consistent gate was defined in the green channel to determine the percentage of peptide-positive cells across all experiments.

### Confocal microscopy

A549 cells were seeded into 24-well plates (50,000 cells/well) and grown for 24 h to approximately 60% confluence. Cells were incubated with 20 µM of FM-labelled peptides (FITC) for 6h at 37°C and the media was then removed. After being washed twice with PBS, pH 7,4, cells were fixed in 2% paraformaldehyde in PBS for 10 min at room temperature. Cells were rinsed twice with PBS and nuclei were stained with 300 nM DAPI in PBS for 10 min at room temperature. The cells were washed again and then analyzed by Nikon-AR1confocal microscope equipped with 3 laser units (405nm for DAPI and 488nm wavelength for FITC). Negative control was performed by omitting the labelled peptide (data not shown).

### Cell Viability Assay

The cytotoxic effect of Cy-9B on A549 cells was evaluated using the MTT (3-(4,5-di-methylthiazol-2-yl)-2,5-diphenyltetrazolium bromide) assay. Equal number of cells (5,000 cells/well) were plated in 96-well plates in triplicate. After 24 hours, attached cells were exposed to increasing concentrations of Cy-9B for 24, 48 and 72 hours. Thereafter, 0.4 mg/mL MTT solution was prepared from stock MTT solution (5 mg/mL PBS) was added per well according to the manufacturer’s protocol an incubated for 4 hours at 37 °C. This was followed by washing with PBS and addition of 100 µL of the DMSO solution into each well to dissolve the formazan crystals. Optical density was recorded at 550 nm. Cell viability (%) was determined by averaging three repeated experiments, with the mean inhibitory concentration (IC50) representing the concentration at which cell viability was reduced by 50%. IC50 were calculated from curves constructed by plotting cell viability (%) versus peptide concentration (µM). Mortality was evaluated by trypan blue staining. A549 cells were exposed to 50, 100, 200 µM Cy-9B treatments, and samples were collected after 24 h and 48 h. At each time point, cells were gently harvested and mixed with trypan blue solution to allow discrimination between viable cells, retaining membrane integrity, and non-viable cells. Dead cells percentages were calculated as the ratio of stained cells to the total number of cells. All measurements were performed in independent biological replicates.

### MTT assay on primary neuronal cultures

Primary hippocampal neurons were prepared from P1-P2 C57bl6 mice as described in ^95^. The plating medium was B27/neurobasal-A (Gibco-Invitrogen) supplemented with 0.5 mM glutamine (Gibco-Invitrogen), 100 U/mL penicillin, and 100 µg/mL streptomycin (Gibco-Invitrogen). At DIV12 20, 50, and 100 µM of Cy-9B was added to the media. After 24h, neuronal viability was assessed using the MTT assay as previously described. Thiazolyl Blue Tetrazolium Bromide (M5655, Merck, USA) was added to cell cultures and incubated for 4h at 37°C. After that, the culture medium was removed, and HCl/ propanol was added. Absorbance was measured at 540 nm by spectrophotometry.

### Boyden Chamber migration and invasion assay

Cell migration was performed using Boyden chambers, with an 8.0 µm pore size (Corning, NY, USA). 30.000 cells/cm^2^ A549 were plated and treated for 24 hours with 25 and 50 µM Cy-9B or the control peptide. Afterward, cells were detached and 0.4 x 10^6^cell/well were suspended in FBS-free media and loaded into the upper chamber, in the absence or presence of 25 and 50 µM Cy-9B or the control peptide. For the invasion assay, 0.4 x 10^6^cell/ well were suspended in FBS-free media and loaded into the upper chamber of a transwell pre-coated with collagen I, following the manufacturer’s protocol (Trevigen Inc., Minneapolis, MN, USA), in the absence or presence of 25 and 50 µM Cy-9B. The lower chamber was filled with a complete medium supplemented with 10% FBS. After 24 h of incubation (37 °C; 5% CO_2_), cells adherent to the underside of the filters were fixed and permeabilized with 70% ethanol, washed with PBS, and stained with 0.25% crystal violet (Merck Life Science, Srl, Milano, Italy). Cells in four random fields at magnification 20× were visualized through an inverted microscope equipped with a digital camera (Leica). Then, crystal violet was dissolved in 100% ethanol, and its absorbance was read at 590 nm in a multiwell plate reader (Tecan InfinitePro Sunrise 200).

### Wound healing assay

For assessing the cell’s migratory ability in a scratch assay, A549 cells (0.3 × 10^4^ cell/cm^2^) were seeded in 6 well plates. After 72 h, cells reached confluence, and the cell monolayer was scratched using a pipette tip through the central axis of the plate. Following one wash with PBS, cell media was replaced with DMEM high glucose without FBS, in the absence (Ctrl) and in the presence of 25 µM and 50 µM of Cy-9B or of the control peptide (scrambled). Migration of the cells into the scratch was digitally documented after 24 and 48 h, and relative migratory activity was calculated measuring the cell-free areas using the ImageJ software. The wound closure areas were visualized under an inverted microscope connected to a digital camera with a 4× magnification.

### Cell Adhesion assay

The assay was performed as previously reported ^96^. In detail, a 48-well plate was coated with 200 µL of collagen (7.5 µg/ml) or Geltrex (ThermoFisher, 0.2 mg/ml), and incubated for 2 hours at room temperature (RT). After PBS washes, plates were blocked by 30 minutes incubation at 37°C in complete cell culture medium. Human lung carcinoma epithelial cells A549, 30.000 cells/cm^2^, were plated and treated for 24 hours with 25 and 50 µM Cy-9B. Afterward, cells were detached, suspended in FBS-free media in the absence or presence of 25 or 50 µM Cy-9B peptide, and 0.4 × 10^6^cell/well were then plated onto the pre-coated wells, and allowed to adhere to the substrates for 30 minutes, at 37°C. Non-adhering cells were washed off with PBS, and the adhering cells were fixed with cold methanol (Sigma-Aldrich) and stained with 0.25% crystal violet. Digital photos of at least 3 random fields were taken, magnification 20×, with an inverted microscope (Leica). The images obtained have been analyzed with the ImageJ software. The data are expressed as the mean percentages ± SEM from at least three independent experiments.

### Primary hippocampal neuronal cultures

Primary neurons were isolated from the dissected hippocampus of P0-P1 mice as previously described in ^97^ with some modifications. Neonatal mice were decapitated, and the brains were transferred to cold Dissection medium (Hank’s Balanced salt solution without calcium and magnesium with 1% of sodium pyruvate 11mg/ml, 0,1% of glucose 20% and 10mM of HEPES 1M). The hippocampi were isolated under a dissecting microscope and were digested in water bath at 37°C for 20 minutes by using two 15-ml tubes, each containing a maximum of 4-5 hippocampi in 5 ml of dissection medium with 0.2% of trypsin. Then, a DNAse solution was added at room temperature for 5 minutes. At the end of incubation, the hippocampi were washed 2 times with the dissection medium and 2 more times with plating medium (Minimum Essential medium with 10% FBS, 0,45% of glucose 20%, 1% of sodium pyruvate 100mM, 1% l-glutamine 200mM,1% Penicillin/Streptomycin 100x), and were dissociated through gentle repeated pipetting by using a 1000 µl tip. The cells were plated in plating medium in 6-mm plates previously coated with 0.05 mg/ml of poly-l-lysine and incubated at 37°C, 5% CO_2_. Two hours after plating, the medium was replaced with 5 ml of maintenance medium (Neurobasal medium with 2% of B-27 supplement, 1% of Glutamax, 1% of P/S). Half of the medium was replaced with fresh medium every 3-4 days until the time of the experiment. Experiments were performed at DIV13.

### Behavioral assay

All behavioral experiments were conducted between Zeitgeber Time 1 (ZT1) and ZT4 in a temperature- and humidity-controlled behavioral chamber (25°C, 60% relative humidity).

### Social behavior assay (Competition for food)

Social interaction assays were conducted using socially experienced male flies, as described previously ^32,88,89^. Briefly, groups of 8 males of the same genotype were anesthetized 24 h prior to testing and housed together in food-containing vials. On the assay day, flies were gently transferred without anaesthesia to empty vials and food-deprived for 90 min. Following deprivation, flies were allowed to acclimate for 2 min in the presence of a food droplet. Social interactions - defined as physical contact, charging behavior, and wing raise - were video-recorded and manually scored over a 2-min observation period.

### Pharmacological treatments

Cy-9B peptide and scrambled peptide were administered at final concentration of 5 µM. Compounds were mixed into Formula 4-24® Instant Drosophila Medium (Carolina Biological Supply) and water to achieve the desired final concentrations, as previously described^32,34^. The vehicle control consisted of water and Formula 4-24® medium only. Compounds were administered for 12 h (overnight). Immediately following treatment, flies were either tested behaviorally.

### Statistical analysis

Statistical analyses were performed using GraphPad Prism. The specific tests used for each dataset are indicated in the corresponding figure legends. Unless otherwise stated, experiments were conducted in at least three independent biological replicates. When data have a normal distribution, an unpaired two-tailed *t*-test was used. One-way analysis of variance (ANOVA) and Dunnett’s and Tukey’s multiple comparison test were applied for comparisons involving more than two groups. *P* values of <0.05 were considered statistically significant. Significance is reported as follows: ns, not significant; *P ≤ 0.05; **P ≤ 0.01; ***P ≤ 0.001.

### Computational studies

The Cy-9B peptide model was built in Chimera ^98^. The protein eIF4E model was obtained from the PDB code 1WKW. The system was then processed using tleap amber tools ^99^. Conventional Cy-9B residues were parametrized using Amber ff14SB forcefield, whereas the two staple components B1K and B2K, were treated as separate residues and parameterized using the generalized Amber forcefield (gaff) ^100^. Water molecules were parametrized using the TIP3P model. The generated coordinates and topology files were then translated into Gromacs input files using the Amber tool parmed ^101^. The potential energy of the complex was then minimized using the gradient descent algorithm. Furthermore, the system was thermalized via a sequence of 14 alternating NPT/NVT steps, increasing the temperature from 50K to 300K while maintaining the pressure at 1 bar during NPT steps. Throughout thermalization, Cα atoms of both protein and peptide were restrained with progressively decreasing force constants. Production simulations were carried out in the NPT ensemble using a v-rescale thermostat and Parrinello-Rahman barostat, with coupling time of 2 ps and 5 ps, respectively. A 2 fs integration timestep was used, and LINCS constraints were applied to all bonds involving hydrogen. For Funnel-Metadynamics (FM), the funnel was positioned using the VMD plug-in funnel.tcl ^102,103^. FM calculations were performed using 10 parallel walkers, each lasting 250 ns, for a total of 2.5 µs of sampling. Equilibrium simulations of the three initial peptide variants were set up following the same workflow, with 500 ns production trajectories. The α-helical content was quantified using an in-house analysis script, as described previously ^21^.

## Acknowledgments

This work is dedicated to Daniele Di Marino, whose vision, guidance, and unwavering dedication inspired this project.

We would like to thank Andrea Frontini for assistance with Confocal Microscopy and Fabio Marcheggiani for Cito-Fluorimetry experiments.

## Funding

This work has received the support of the Italian Foundation for Cancer Research AIRC (Project No. IG 807 2022 ID 27534) and supported in part by a grant from FRAXA Research Foundation. This work has received funding from the European Research Council (ERC) under the European Union’s Horizon 2020 research and innovation programme (“CoMMBi” ERC grant agreement No.101001784), the Swiss National Science Foundation (SNSF) (grant number IC00I0-231546) and a grant from the Swiss National Super-computing Centre (CSCS) under project ID u8. GV was supported by the Istituto Superiore di Sanità-UNIVPM doctoral program.

## Author contributions

Conceptualization: ARm, DDM

Methodology: ARm, SR, AG, ADL, TB

Investigation: ARm, SR, OA, AG, ADL, JS, RP, ZB,GV

Visualization: JR, ARs, GB

Supervision: ARm, DDM, VL

Writing—original draft: ARm, SR, VL

Writing—review & editing: ALT, CB, AM, JR, AC, VL

ARm: Alice Romagnoli

ARs: Agnese Roscioni

## Competing interests

Authors declare that they have no competing interests.

## Data and materials availability

All data are available in the main text or the supplementary materials.

## Notes

### Competing Interest Statement

The authors have declared no competing interest.

## References

1. Costa-Mattioli, M., Sossin, W. S., Klann, E. & Sonenberg, N. Translational Control of Long-Lasting Synaptic Plasticity and Memory. Neuron 61, 10–26 (2009).

2. Bhat, M. et al. Targeting the translation machinery in cancer. Nat. Rev. Drug Discov. 14, 261–278 (2015).

3. Sonenberg, N. & Hinnebusch, A. G. Regulation of Translation Initiation in Eukaryotes: Mechanisms and Biological Targets. Cell 136, 731–745 (2009).

4. Merrick, W. C. & Pavitt, G. D. Protein Synthesis Initiation in Eukaryotic Cells. Cold Spring Harb. Perspect. Biol. 10, a033092 (2018).

5. Romagnoli, A. et al. Control of the eIF4E activity: structural insights and pharmacological implications. Cellular and Molecular Life Sciences 78, 6869–6885 (2021).

6. De Benedetti, A. & Graff, J. R. eIF-4E expression and its role in malignancies and metastases. Oncogene 23, 3189–3199 (2004).

7. Siddiqui, N. & Sonenberg, N. Signalling to eIF4E in cancer. Biochem. Soc. Trans. 43, 763–772 (2015).

8. Dai, L. et al. Targeting EIF4F complex in non-small cell lung cancer cells. Oncotarget 8, 55731–55735 (2017).

9. Tan, H., He, L. & Cheng, Z. Inhibition of eIF4E signaling by ribavirin selectively targets lung cancer and angiogenesis. Biochem. Biophys. Res. Commun. 529, 519–525 (2020).

10. Dela Cruz, C. S., Tanoue, L. T. & Matthay, R. A. Lung Cancer: Epidemiology, Etiology, and Prevention. Clin. Chest Med. 32, 605–644 (2011).

11. Richter, J. D. & Zhao, X. The molecular biology of FMRP: new insights into fragile X syndrome. Nat. Rev. Neurosci. 22, 209–222 (2021).

12. Romagnoli, A. & Di Marino, D. The Use of Peptides in the Treatment of Fragile X Syndrome: Challenges and Opportunities. Front. Psychiatry 12, (2021).

13. O’Rourke, R. L. & Garner, A. L. Chemical Probes for Studying the Eukaryotic Translation Initiation Factor 4E (eIF4E)-Regulated Translatome in Cancer. ACS Pharmacol. Transl. Sci. 8, 621–635 (2025).

14. Maracci, C. et al. The mTOR/4E-BP1/eIF4E Signalling Pathway as a Source of Cancer Drug Targets. Curr. Med. Chem. 29, 3501–3529 (2022).

15. Romagnoli, A., Maracci, C., D’Agostino, M., La Teana, A. & Di Marino, D. Targeting mTOR and eIF4E: a feasible scenario in ovarian cancer therapy. Cancer Drug Resistance 10.20517/cdr.2021.20(2021) doi:10.20517/cdr.2021.20.

16. Lu, C., Makala, L., Wu, D. & Cai, Y. Targeting translation: eIF4E as an emerging anticancer drug target. Expert Rev. Mol. Med. 18, e2 (2016).

17. Roscioni, A. et al. Ligand-Induced Structural Dynamics Drive Allosteric Regulation of Translation Initiation Factor eIF4E. Preprint at 10.64898/2025.12.23.696204 (2025).

18. Peter, D. et al. Molecular Architecture of 4E-BP Translational Inhibitors Bound to eIF4E. Mol. Cell 57, 1074–1087 (2015).

19. Marcotrigiano, J., Gingras, A.-C., Sonenberg, N. & Burley, S. K. Cap-Dependent Translation Initiation in Eukaryotes Is Regulated by a Molecular Mimic of eIF4G. Mol. Cell 3, 707–716 (1999).

20. Di Marino, D. et al. MD and Docking Studies Reveal That the Functional Switch of CYFIP1 is Mediated by a Butterfly-like Motion. J. Chem. Theory Comput. 11, 3401–3410 (2015).

21. Di Marino, D., D’Annessa, I., Tancredi, H., Bagni, C. & Gallicchio, E. A unique binding mode of the eukaryotic translation initiation factor 4E for guiding the design of novel peptide inhibitors. Protein Science 24, 1370–1382 (2015).

22. Grüner, S. et al. The Structures of eIF4E-eIF4G Complexes Reveal an Extended Interface to Regulate Translation Initiation. Mol. Cell 64, 467–479 (2016).

23. Napoli, I. et al. The Fragile X Syndrome Protein Represses Activity-Dependent Translation through CYFIP1, a New 4E-BP. Cell 134, 1042–1054 (2008).

24. De Rubeis, S. et al. CYFIP1 Coordinates mRNA Translation and Cytoskeleton Remodeling to Ensure Proper Dendritic Spine Formation. Neuron 79, 1169–1182 (2013).

25. Verkerk, A. J. M. H. et al. Identification of a gene (FMR-1) containing a CGG repeat coincident with a breakpoint cluster region exhibiting length variation in fragile X syndrome. Cell 65, 905–914 (1991).

26. D’Annessa, I., Cicconardi, F. & Di Marino, D. Handling FMRP and its molecular partners: Structural insights into Fragile X Syndrome. Prog. Biophys. Mol. Biol. 141, 3–14 (2019).

27. Bagni, C., Tassone, F., Neri, G. & Hagerman, R. Fragile X syndrome: causes, diagnosis, mechanisms, and therapeutics. Journal of Clinical Investigation 122, 4314–4322 (2012).

28. Chen, Z. et al. Structure and control of the actin regulatory WAVE complex. Nature 468, 533–538 (2010).

29. Han, K. A. & Ko, J. Orchestration of synaptic functions by WAVE regulatory complex-mediated actin reorganization. Exp. Mol. Med. 55, 1065–1075 (2023).

30. DomínguezIturza, N. et al. The autism- and schizophrenia-associated protein CYFIP1 regulates bilateral brain connectivity and behaviour. Nat. Commun. 10, 3454 (2019).

31. Woo, Y. J. et al. Domain-Specific Cognitive Impairments in Humans and Flies With Reduced CYFIP1 Dosage. Biol. Psychiatry 86, 306–314 (2019).

32. Kanellopoulos, A. K. et al. Aralar Sequesters GABA into Hyperactive Mitochondria, Causing Social Behavior Deficits. Cell 180, 1178–1197.e20 (2020).

33. DomínguezIturza, N. et al. The autism- and schizophrenia-associated protein CYFIP1 regulates bilateral brain connectivity and behaviour. Nat. Commun. 10, 3454 (2019).

34. Mariano, V. et al. SREBP modulates the NADP+/NADPH cycle to control night sleep in Drosophila. Nat. Commun. 14, 763 (2023).

35. Teng, Y. et al. The WASF3–NCKAP1–CYFIP1 Complex Is Essential for Breast Cancer Metastasis. Cancer Res. 76, 5133–5142 (2016).

36. Teng, Y. et al. Targeting the WASF3–CYFIP1 Complex Using Stapled Peptides Suppresses Cancer Cell Invasion. Cancer Res. 76, 965–973 (2016).

37. Moerke, N. J. et al. Small-Molecule Inhibition of the Interaction between the Translation Initiation Factors eIF4E and eIF4G. Cell 128, 257–267 (2007).

38. Papadopoulos, E. et al. Structure of the eukaryotic translation initiation factor eIF4E in complex with 4EGI-1 reveals an allosteric mechanism for dissociating eIF4G. Proceedings of the National Academy of Sciences 111, (2014).

39. Ko, S. Y., Guo, H., Barengo, N. & Naora, H. Inhibition of Ovarian Cancer Growth by a Tumor-Targeting Peptide That Binds Eukaryotic Translation Initiation Factor 4E. Clinical Cancer Research 15, 4336–4347 (2009).

40. Lama, D., Quah, S. T., Brown, C. J., Lane, D. P. & Verma, C. S. 159 Stapled-peptides targeting the protein-binding interface of eukaryotic Translation Initiation Factor 4E (eIF4E) protein. J. Biomol. Struct. Dyn. 33, 102–103 (2015).

41. Masse, M. et al. An eIF4E-interacting peptide induces cell death in cancer cell lines. Cell Death Dis. 5, e1500–e1500 (2014).

42. Gallagher, E. E. et al. A cell-penetrant lactam-stapled peptide for targeting eIF4E protein-protein interactions. Eur. J. Med. Chem. 205, 112655 (2020).

43. Fischer, P. D. et al. A biphenyl inhibitor of eIF4E targeting an internal binding site enables the design of cell-permeable PROTAC-degraders. Eur. J. Med. Chem. 219, 113435 (2021).

44. Frosi, Y. et al. Engineering an autonomous VH domain to modulate intracellular pathways and to interrogate the eIF4F complex. Nat. Commun. 13, 4854 (2022).

45. Qi, X. et al. EGPI-1, a novel eIF4E/eIF4G interaction inhibitor, inhibits lung cancer cell growth and angiogenesis through Ras/MNK/ERK/eIF4E signaling pathway. Chem. Biol. Interact. 352, 109773 (2022).

46. Chen, B. et al. The WAVE Regulatory Complex Links Diverse Receptors to the Actin Cytoskeleton. Cell 156, 195–207 (2014).

47. Fagiani, F. et al. Pin1 as Molecular Switch in Vascular Endothelium: Notes on Its Putative Role in Age-Associated Vascular Diseases. Cells 10, 3287 (2021).

48. Wang, H. et al. Peptide-based inhibitors of protein–protein interactions: biophysical, structural and cellular consequences of introducing a constraint. Chem. Sci. 12, 5977–5993 (2021).

49. Lombardi, L., Genio, V. Del, Albericio, F. & Williams, D. R. Advances in Peptidomimetics for Next-Generation Therapeutics: Strategies, Modifications, and Applications. Chem. Rev. 10.1021/acs.chemrev.4c00989 (2025) doi:10.1021/acs.chemrev.4c00989.

50. Ali, A. M., Atmaj, J., Van Oosterwijk, N., Groves, M. R. & Dömling, A. Stapled Peptides Inhibitors: A New Window for Target Drug Discovery. Comput. Struct. Biotechnol. J. 17, 263–281 (2019).

51. D’Annessa, I. et al. Bioinformatics and Biosimulations as Toolbox for Peptides and Peptidomimetics De-sign: Where Are We? Front. Mol. Biosci. 7, (2020).

52. Qvit, N., Rubin, S. J. S., Urban, T. J., Mochly-Rosen, D. & Gross, E. R. Peptidomimetic therapeutics: scientific approaches and opportunities. Drug Discov. Today 22, 454–462 (2017).

53. Limongelli, V., Bonomi, M. & Parrinello, M. Funnel metadynamics as accurate binding free-energy method. Proceedings of the National Academy of Sciences 110, 6358–6363 (2013).

54. Baar, S., Kuragano, M., Tokuraku, K. & Watanabe, S. Towards a comprehensive approach for characterizing cell activity in bright-field microscopic images. Sci. Rep. 12, 16884 (2022).

55. Zhu, J., Wang, Y., Gao, F. & Zhang, Z. Optical differentiation in a polar coordinate system. Appl. Phys. Lett. 122, (2023).

56. Tomoo, K. et al. Structural basis for mRNA Cap-Binding regulation of eukaryotic initiation factor 4E by 4E-binding protein, studied by spectroscopic, X-ray crystal structural, and molecular dynamics simulation methods. Biochimica et Biophysica Acta (BBA) - Proteins and Proteomics 1753, 191–208 (2005).

57. Brown, C. J., McNae, I., Fischer, P. M. & Walkinshaw, M. D. Crystallographic and Mass Spectrometric Characterisation of eIF4E with N7-alkylated Cap Derivatives. J. Mol. Biol. 372, 7–15 (2007).

58. Brown, C. J., Verma, C. S., Walkinshaw, M. D. & Lane, D. P. Crystallization of eIF4E complexed with eIF4GI peptide and glycerol reveals distinct structural differences around the cap-binding site. Cell Cycle 8, 1905–1911 (2009).

59. Fukuyo, A., In, Y., Ishida, T. & Tomoo, K. Structural scaffold for eIF4E binding selectivity of 4E-BP isoforms: crystal structure of eIF4E binding region of 4E-BP2 and its comparison with that of 4E-BP1. Journal of Peptide Science 17, 650–657 (2011).

60. Siddiqui, N. et al. Structural Insights into the Allosteric Effects of 4EBP1 on the Eukaryotic Translation Initiation Factor eIF4E. J. Mol. Biol. 415, 781–792 (2012).

61. Di Marino, D. et al. Binding of the Anti-FIV Peptide C8 to Differently Charged Membrane Models: From First Docking to Membrane Tubulation. Front. Chem. 8, (2020).

62. Zantza, I. et al. Uracil/H+ Symport by FurE Refines Aspects of the Rocking-bundle Mechanism of APC-type Transporters. J. Mol. Biol. 435, 168226 (2023).

63. Moraca, F. et al. Ligand binding to telomeric G-quadruplex DNA investigated by funnel-metadynamics simulations. Proceedings of the National Academy of Sciences 114, (2017).

64. Comitani, F., Limongelli, V. & Molteni, C. The Free Energy Landscape of GABA Binding to a Pentameric Ligand-Gated Ion Channel and Its Disruption by Mutations. J. Chem. Theory Comput. 12, 3398–3406 (2016).

65. Yuan, X., Raniolo, S., Limongelli, V. & Xu, Y. The Molecular Mechanism Underlying Ligand Binding to the Membrane-Embedded Site of a G-Protein-Coupled Receptor. J. Chem. Theory Comput. 14, 2761–2770 (2018).

66. Troussicot, L., Guillière, F., Limongelli, V., Walker, O. & Lancelin, J.-M. Funnel-Metadynamics and Solution NMR to Estimate Protein–Ligand Affinities. J. Am. Chem. Soc. 137, 1273–1281 (2015).

67. Brotzakis, Z. F., Limongelli, V. & Parrinello, M. Accelerating the Calculation of Protein–Ligand Binding Free Energy and Residence Times Using Dynamically Optimized Collective Variables. J. Chem. Theory Comput. 15, 743–750 (2019).

68. Zhou, W. et al. Improved eIF4E Binding Peptides by Phage Display Guided Design: Plasticity of Interacting Surfaces Yield Collective Effects. PLoS One 7, e47235 (2012).

69. Lama, D. et al. Rational Optimization of Conformational Effects Induced By Hydrocarbon Staples in Peptides and their Binding Interfaces. Sci. Rep. 3, 3451 (2013).

70. Penchala, S. C. et al. A biomimetic approach for enhancing the in vivo half-life of peptides. Nat. Chem. Biol. 11, 793–798 (2015).

71. Lombardi, L., Genio, V. Del, Albericio, F. & Williams, D. R. Advances in Peptidomimetics for Next-Generation Therapeutics: Strategies, Modifications, and Applications. Chem. Rev. 10.1021/acs.chemrev.4c00989 (2025) doi:10.1021/acs.chemrev.4c00989.

72. Romagnoli, A. et al. Peptidomimetics design and characterization: Bridging experimental and computer-based approaches. in 279–327 (2025). doi:10.1016/bs.pmbts.2024.07.002.

73. Peng, X., Wellard, N., Ghosh, A., Troakes, C. & Giese, K. P. Different dysregulations of CYFIP1 and CYFIP2 in distinct types of dementia. Brain Res. Bull. 206, 110849 (2024).

74. Dowling, R. J. O. et al. mTORC1-Mediated Cell Proliferation, But Not Cell Growth, Controlled by the 4E-BPs. Science (1979). 328, 1172–1176 (2010).

75. Woodcock, H. V. et al. The mTORC1/4E-BP1 axis represents a critical signaling node during fibrogenesis. Nat. Commun. 10, 6 (2019).

76. Gandin, V. et al. Cap-dependent translation initiation monitored in living cells. Nat. Commun. 13, 6558 (2022).

77. Fabbri, L., Chakraborty, A., Robert, C. & Vagner, S. The plasticity of mRNA translation during cancer progression and therapy resistance. Nat. Rev. Cancer 21, 558–577 (2021).

78. Pettersson, F. et al. Genetic and Pharmacologic Inhibition of eIF4E Reduces Breast Cancer Cell Migration, Invasion, and Metastasis. Cancer Res. 75, 1102–1112 (2015).

79. Jacquemont, S. et al. Protein synthesis levels are increased in a subset of individuals with fragile X syndrome. Hum. Mol. Genet. 27, 2039–2051 (2018).

80. Berry-Kravis, E. et al. A Double-Blind, Randomized, Placebo-Controlled Clinical Study of Trofinetide in the Treatment of Fragile X Syndrome. Pediatr. Neurol. 110, 30–41 (2020).

81. Santini, E. et al. Reducing eIF4E-eIF4G interactions restores the balance between protein synthesis and actin dynamics in fragile X syndrome model mice. Sci. Signal. 10, (2017).

82. Kleczewska, N. et al. Cellular delivery of dinucleotides by conjugation with small molecules: targeting translation initiation for anticancer applications. Chem. Sci. 12, 10242–10251 (2021).

83. Hagerman, R. J. et al. Fragile X syndrome. Nat. Rev. Dis. Primers 3, 17065 (2017).

84. Gillett, D. A., Tigro, H., Wang, Y. & Suo, Z. FMR1 Disorders: Basics of Biology and Therapeutics in Development. Cells 13, 2100 (2024).

85. Shah, S. et al. Antisense oligonucleotide rescue of CGG expansion–dependent *FMR1* mis-splicing in fragile X syndrome restores FMRP. Proceedings of the National Academy of Sciences 120, (2023).

86. Berry-Kravis, E. M. et al. Inhibition of phosphodiesterase-4D in adults with fragile X syndrome: a randomized, placebo-controlled, phase 2 clinical trial. Nat. Med. 27, 862–870 (2021).

87. Berry-Kravis, E. et al. A randomized, controlled trial of ZYN002 cannabidiol transdermal gel in children and adolescents with fragile X syndrome (CONNECT-FX). J. Neurodev. Disord. 14, 56 (2022).

88. Edwards, A. C., Rollmann, S. M., Morgan, T. J. & Mackay, T. F. C. Quantitative Genomics of Aggressive Behavior in Drosophila melanogaster. PLoS Genet. 2, e154 (2006).

89. Rollmann, S. M. et al. Pleiotropic Effects of Drosophila *neuralized* on Complex Behaviors and Brain Structure. Genetics 179, 1327–1336 (2008).

90. Zhang, Y. et al. Improving Stability Enhances In Vivo Efficacy of a PCSK9 Inhibitory Peptide. J. Am. Chem. Soc. 144, 19485–19498 (2022).

91. Bruni, R. & Kloss, B. High-Throughput Cloning and Expression of Integral Membrane Proteins in *Escherichia coli*. Curr. Protoc. Protein Sci. 74, (2013).

92. Schmidt, E. K., Clavarino, G., Ceppi, M. & Pierre, P. SUnSET, a nonradioactive method to monitor protein synthesis. Nat. Methods 6, 275–277 (2009).

93. Niesner U. et al. Quantitation of the Tumor-Targeting Properties of Antibody Fragments Conjugated to Cell-Permeating HIV-1 TAT Peptides. vol. 13 729–736 Preprint at 10.1021/bc025517 (2002).

94. Krämer, S. D. & Wunderli-Allenspach, H. No entry for TAT(44–57) into liposomes and intact MDCK cells: novel approach to study membrane permeation of cell-penetrating peptides. Biochimica et Biophysica Acta (BBA) - Biomembranes 1609, 161–169 (2003).

95. Sclip, A. et al. Soluble Aβ oligomer-induced synaptopathy: c-Jun N-terminal kinase’s role. J. Mol. Cell Biol. 5, 277–279 (2013).

96. Luca, A. De et al. The nitrobenzoxadiazole derivative MC3181 blocks melanoma invasion and metastasis. Oncotarget 8, 15520–15538 (2017).

97. Beaudoin, G. M. J. et al. Culturing pyramidal neurons from the early postnatal mouse hippocampus and cortex. Nat. Protoc. 7, 1741–1754 (2012).

98. Pettersen, E. F. et al. UCSF Chimera—A visualization system for exploratory research and analysis. J. Comput. Chem. 25, 1605–1612 (2004).

99. Case, D. A. et al. AmberTools. J. Chem. Inf. Model. 63, 6183–6191 (2023).

100. Maier, J. A. et al. ff14SB: Improving the Accuracy of Protein Side Chain and Backbone Parameters from ff99SB. J. Chem. Theory Comput. 11, 3696–3713 (2015).

101. Abraham, M. J. et al. GROMACS: High performance molecular simulations through multi-level parallelism from laptops to supercomputers. SoftwareX 1–2, 19–25 (2015).

102. Raniolo, S. & Limongelli, V. Ligand binding free-energy calculations with funnel metadynamics. Nat. Protoc. 15, 2837–2866 (2020).

103. Humphrey, W., Dalke, A. & Schulten, K. VMD: Visual molecular dynamics. J. Mol. Graph. 14, 33–38 (1996).

