## Supplementary material for "A CYFIP1-Inspired Peptidomimetic Modulates eIF4E-Dependent Translational Control in Cancer and Neurodevelopmental Disorders": Supplemetary Information

**This PDF file includes:**

Figs. S1 to S8
Table S1

**Other Supplementary Materials for this manuscript include the following:**

Video S1

| | | |
|---|---|---|
| **eIF4G1-iso 1** | 605-NLEEKKRYDREFLLGFQFIFAS | canonical |
| **eIF4G1-iso 3** | 565-NLEEKKRYDREFLLGFQFIFAS | alternative initiation Met-41 |
| **eIF4G1-iso 4** | 518-NLEEKKRYDREFLLGFQFIFAS | alternative initiation Met-88 |
| **eIF4G1-iso 5** | 441-NLEEKKRYDREFLLGFQFIFAS | alternative initiation Met-165 |
| **eIF4G1-iso 6** | 409-NLEEKKRYDREFLLGFQFIFAS | alternative initiation Met-197 |
| **eIF4G1-iso 7** | 409-NLEEKKRYDREFLLGFQFIFAS | alternative splicing |
| **eIF4G1-iso 8** | 605-NLEEKKRYDREFLLGFQFIFAS | alternative splicing |
| **eIF4G1-iso 9** | 612-NLEEKKRYDREFLLGFQFIFAS | alternative splicing |
| **eIF4G3-iso 1** | 617-DTEGKKQYDREFLLDFQFMPAC | canonical |
| **eIF4G3-iso 3** | 623-DTEGKKQYDREFLLDFQFMPAC | alternative splicing |
| **eIF4G3-iso 4** | 337-DTEGKKQYDREFLLDFQFMPAC | alternative splicing |
| **4EBP1** | 47-PGGTRIIYDRKFLMECRNSPVT | |
| **4EBP2** | 47-PGGTRIIYDRKFLLDRRNSPMA | |
| **4EBP3** | 33-PGGTRIIYDRKFLLECKNSPIA | |
| **4T-iso 1** | 23-ASKCPHRYTKEELLDIKELPHS | canonical |
| **4T-iso 2** | 23-ASKCPHRYTKEELLDIKELPHS | alternative splicing |
| **4T-iso 3** | 23-ASKCPHRYTKEELLDIKELPHS | alternative splicing |
| **CYFIP1-iso 1** | 716-VMAGSLLLDKRLRSECKNQGAT | canonical |
| **CYFIP1-iso 2** | 285-VMAGSLLLDKRLRSECKNQGAT | alternative splicing |
| **CYFIP2-iso 1** | 740-AMAGSVLLDKRFRAECKNYGVI | canonical |
| **CYFIP2-iso 2** | 715-AMAGSVLLDKRFRAECKNYGVI | alternative splicing |

Non-polar
Polar
Positive charge
Negative charge

**Fig. S1. Multiple sequence alignment of eIF4E-interacting regions across 4E-BP family members.** The first column lists the proteins and isoforms analyzed; the second column displays the aligned sequences; and the third column indicates whether each corresponds to a canonical or alternative spliced variant. Residues within the canonical eIF4E-binding site are color-coded by biochemical class: non-polar (green), polar (light blue), positively charged (yellow), and negatively charged (red).

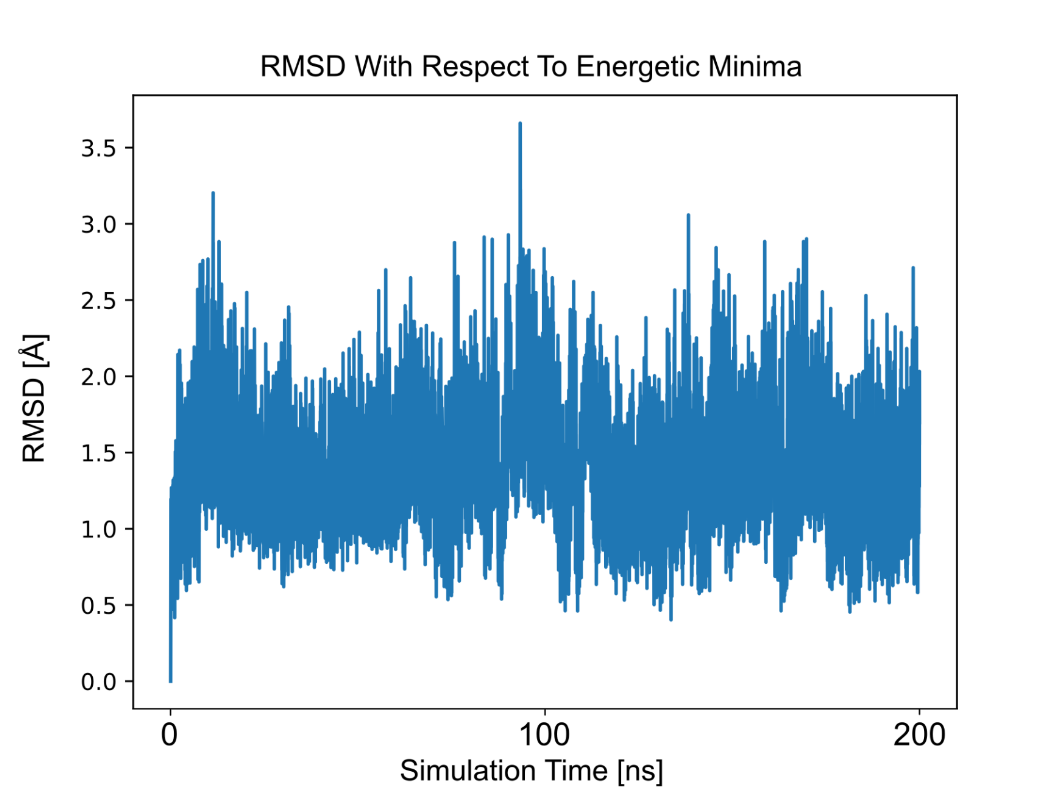


**Fig. S2. Unbiased simulation of the Cy-9B–eIF4E complex at the predicted lowest energy minimum.** The RMSD values from a 200ns unbiased molecular dynamics simulation are shown for Cy-9B bound to eIF4E. The starting state of the simulation is the binding mode identified as the lowest energy minimum by FM calculations. RMSD was calculated on the Cα atoms of the helical region of Cy-9B to monitor the stability of the predicted binding mode.

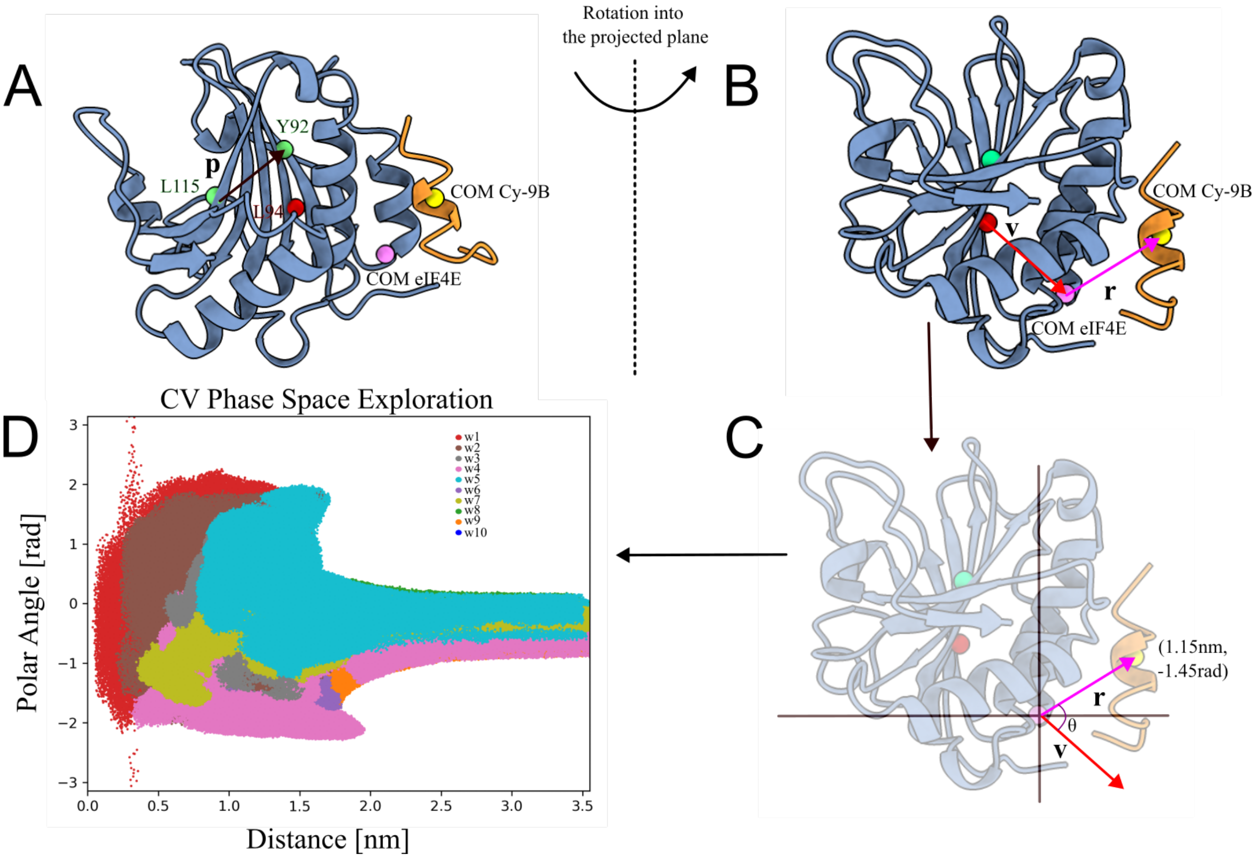


**Fig. S3. Definition of polar-coordinate collective variables (CVs). (A)** Illustration of the projection vector *p*, which defines the 2D plane used to construct the polar-coordinate CV system. The vector *p* is obtained from the coordinates of the Cα atoms of residues L115 and Y92 of eIF4E (green spheres). **(B)** Geometric representation of the polar-coordinate CVs employed in the FM calculations, where vector *p* is perpendicular to the viewer. Within this plane, two vectors, *v* and *r*, are defined to establish the polar CVs. Vector *v* is constructed between the Cα atom of residue L94 (red sphere) and the center of mass (COM) of selected eIF4E pocket residues (pink sphere). Vector *r* spans from the COM of the same eIF4E pocket residues to the COM of the Cα atoms of the Cy-9B helical region (yellow sphere). **(C)** Schematic of the resulting polar-coordinate CV space. The radial CV (*r*) measures the distance between the COM of the eIF4E binding pocket and the Cy-9B helical COM. The angular CV ($\theta$) is defined as the angle between vectors *r* and *v*. Example values for the illustrated pose are shown (1.15 nm and -1.45 radians). **(D)** Sampling distribution across the 2D CV space (distance and polar angle) during the production FM run. Data from all 10 multiple walkers are overlaid and color-coded to visualize exploration of the CV landscape.

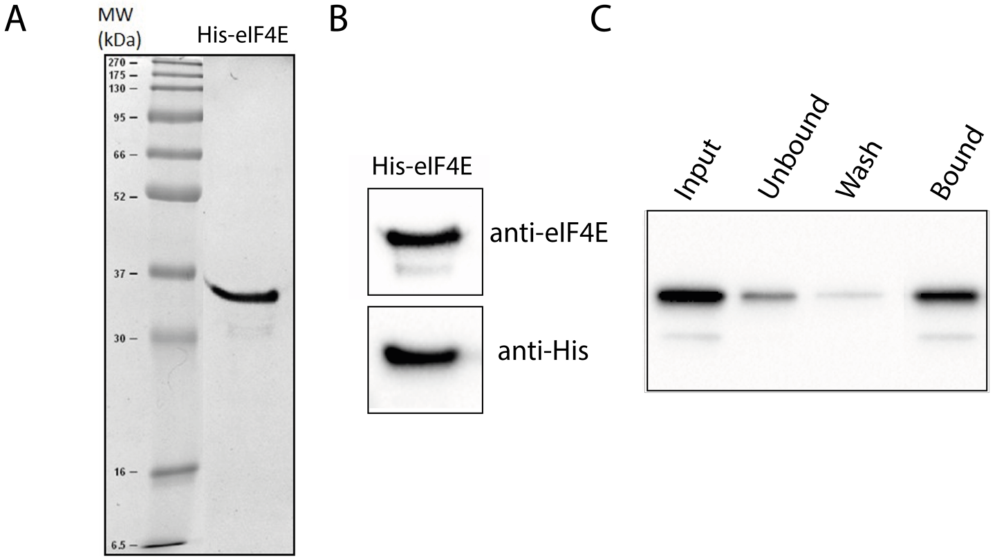


**Fig. S4. Production and activity assay of recombinant N-His-eIF4E. (A)** Purification of N-His-eIF4E expressed in *E. coli* BL21 (DE3) pLysS, visualized on a Coomassie-stained 12% SDS- PAGE gel. (**B)** Western blot validation of purified protein using anti-eIF4E and anti-histidine antibodies. (**C)** $m^7$GTP pull-down assay performed with recombinant eIF4E. Western blot analysis showing the different steps of the pull-down process, with immunodetection performed using anti-eIF4E antibodies.

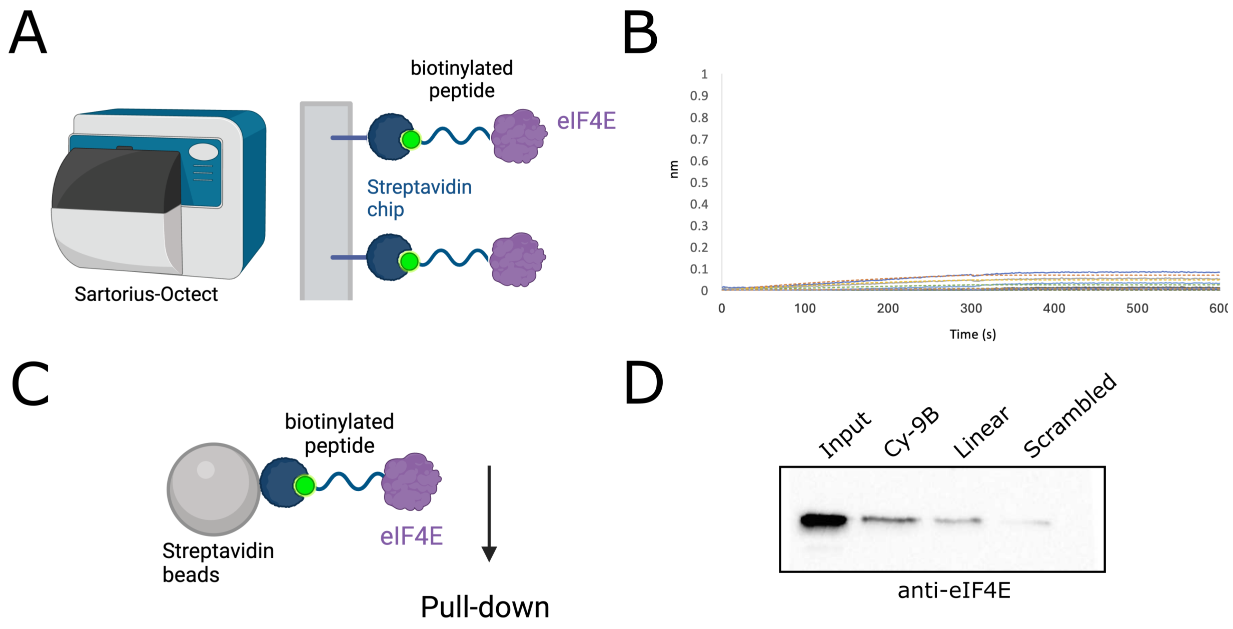


**Fig. S5. *In vitro* binding analysis of eIF4E with Cy-9B and control peptides. (A)** Schematic of Bio-Layer Interferometry (BLI) assay and (**B**) representative sensorgram for the scrambled control peptide. **(C)** Schematic of the streptavidin magnetic beads pull-down assay using biotinylated Cy-9B, scrambled, or linear peptides immobilized on streptavidin-coated beads. (**D**) Bound fractions were analyzed by Western blot using anti-eIF4E antibody.

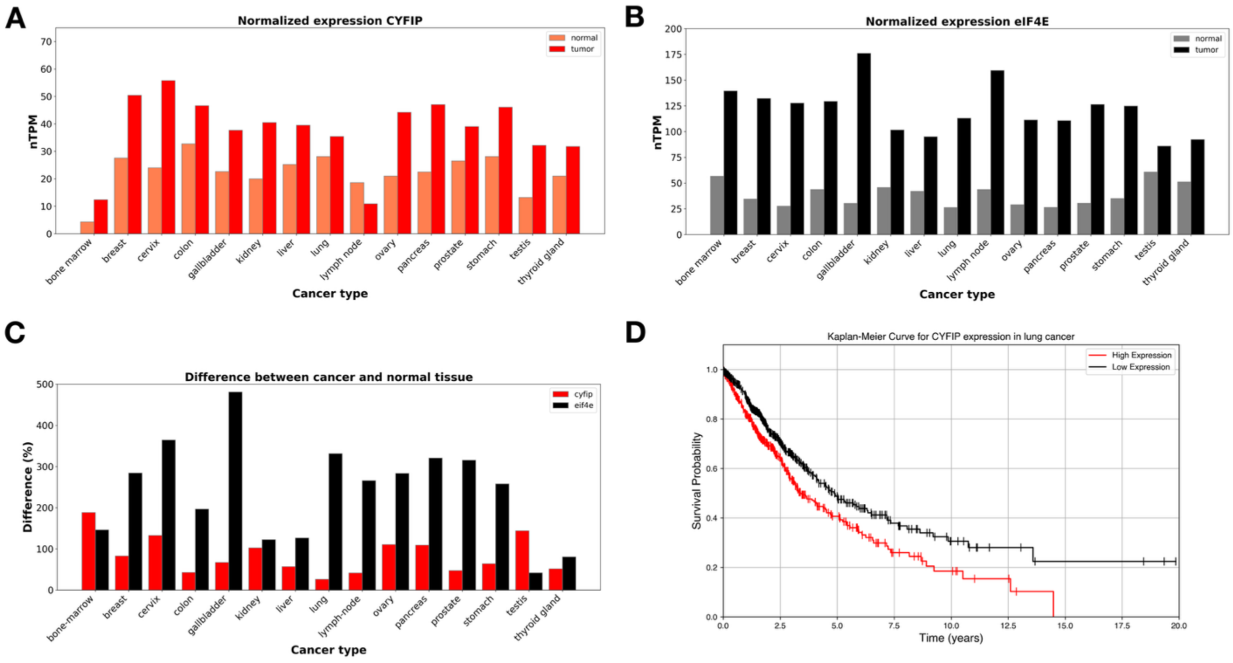


**Fig. S6. Expression levels of eIF4E and CYFIP1 across cancer types. (A-B)** Normalized transcript levels per million (nTMP) of CYFIP1 **(A)** and eIF4E **(B)** in normal and cancer cells from multiple tissues. **(C)** Percentage difference in expression between cancer and matched normal tissues for CYFIP1 (red) and eIF4E (black) across multiple tissue types. **(D)** Kaplan-Meier survival curve of CYFIP1 expression in lung cancer calculated over 994 patients, showing the association between CYFIP1 levels and mortality. Data were obtained from The Human Protein Atlas (version 23.0) and Ensembl (version 109) and normalized using the trimmed mean of M-values.

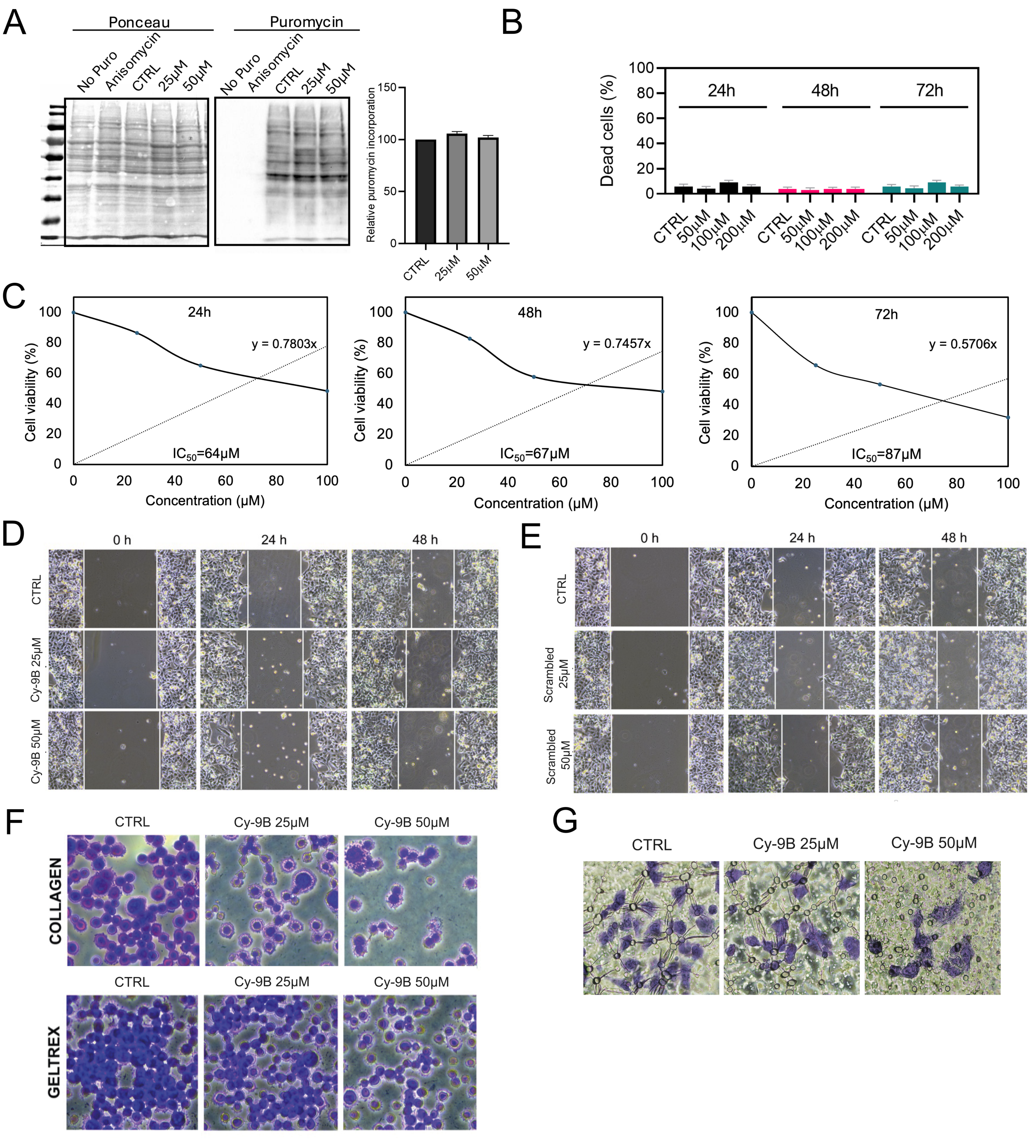


**Fig. S7. Peptide effects on tumor cell translation, viability, adhesion, migration, and invasion.**
**(A)** Western blot analysis of protein synthesis rates assessed using the SUnSET assay after 24-hour treatment with scrambled peptide at different concentrations in A549 cells. Quantification of puromycin incorporation is shown. Ponceau staining was used for normalization. The results are expressed as the fold change relative to wild-type controls (n = 3). Data are mean ± SEM. Statistical significance was determined by one-way ANOVA. (**B)** Quantification of cell death in A549 cells treated with increasing concentrations of Cy-9B after Trypan Blue staining, expressed as the percentage of dead cells. **(C)** Dose-

response curves and $IC_{50}$ values of Cy-9B in A549 cells, determined by MTT assay after 24, 48, and 72 hours of treatment. Cell viability (%) was assessed at increasing peptide concentrations (25, 50 or 100 μM), across three independent experiments. $IC_{50}$ values were calculated as the concentration at which Cy-9B reduced cell viability by 50%, calculated from the fitted curves. **(D-E)** Scratch wound healing assay showing cell motility of A549 cells treated with Cy-9B **(D)** or scrambled peptide **(E)** at 25 and 50 μM for up to 48 hours. A scratch was introduced in the cell monolayer, and wound closure was monitored by phase contrast microscopy at 4× magnification. **(F)** Effect of Cy-9B on A549 cells adhesion to extracellular matrix (ECM) components. Cells ($4x10^5$/ml) were plated to individual coated wells with 200 μl of collagen (7.5 μg/ml), or geltrex (0.2 mg/ml), in the absence or presence of 25 or 50 μM Cy-9B and incubated for 30 minutes at 37°C in 5% $CO_2$. Cells were then fixed and stained with crystal violet. Images were obtained by phase contrast microscopy (20X magnification). **(G)** Invasion assay of A549 cells treated with Cy-9B. A549 cells were pre-treated for 24 hours with 25 and 50 μM Cy-9B. Then cells were plated in the upper part of a Boyder chamber coated with collagen I, in the presence or absence of 25 and 50 μM Cy-9B. After 24 hours invaded cells were stained through Crystal violet, the staining was solubilized with 100% EtOH and quantified by absorbance at 590 nm.

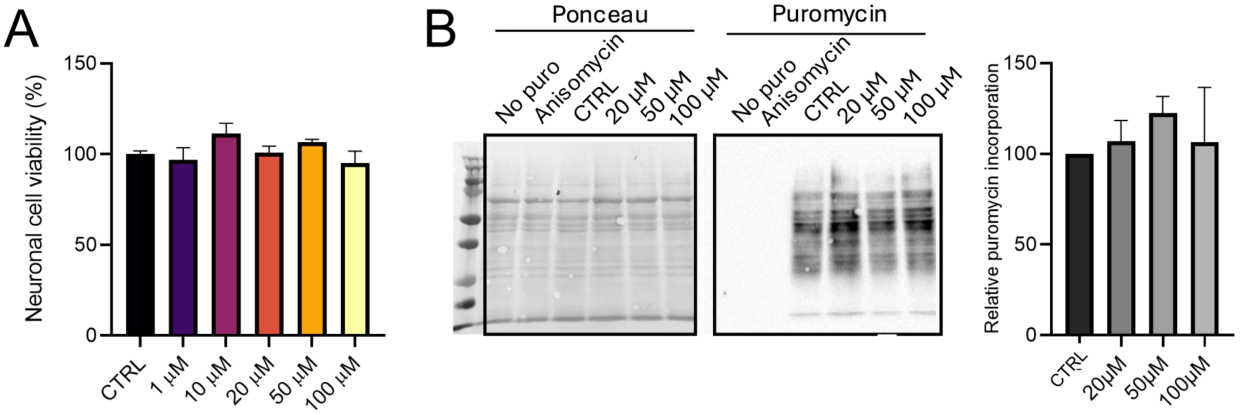


**Figure S8. Effect of peptide on neuronal protein synthesis and viability.**
(**A**) Viability of primary hippocampal neurons assessed by MTT assay after 24 h exposure to Cy-9B (0 to 100 µM). (**B**) Quantitative analysis of puromycin incorporation in *Fmr1*-KO neurons after treatment with different concentrations of scrambled peptide. Ponceau was used for normalization. The results are expressed as the fold change relative to wild-type controls (n = 3). Data are presented as mean ± SEM. Statistical significance is indicated and was determined by one-way ANOVA.

| Peptide name | Sequence | MS calc. | MS found |
|---|---|---|---|
| Cy-lin | Ac-LLLDKRLRSECKNQ-$NH_2$ | [M+1]$^{+}$=1758.07;<br>[M+2]$^{2+}$=879.5;<br>[M+3]$^{3+}$=586.7.1 | [M+2]$^{2+}$=879.4;<br>[M+3]$^{3+}$=586.2 |
| Biot-linear | Biot-[$O_2$Oc]-LLLDKRLRSECKNQ-$NH_2$ | [M+1]$^{+}$=2086.1;<br>[M+2]$^{2+}$=1043.5;<br>[M+3]$^{3+}$=696.0 | [M+2]2+=1043.3;<br>[M+3]3+=695.9 |
| Cy-9B macrocyclic stapled | Ac-LLLDKRLR-$X_1$-ECK-$X_2$-Q-$NH_2$<br>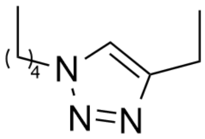 | [M+1]$^{+}$=1805.03;<br>[M+2]$^{2+}$=903.2;<br>[M+3]$^{3+}$=602.5 | [M+2]$^{2+}$=903.7;<br>[M+3]$^{3+}$=602.8 |
| Biot-Cy-9B | Biot-[$O_2$Oc]-LLLDKRLR-$X_1$-ECK-$X_2$-Q-$NH_2$<br>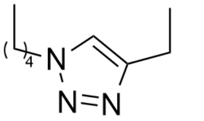 | [M+1]$^{+}$=2134.1;<br>[M+2]$^{2+}$=1067.8;<br>[M+3]$^{3+}$=712.1 | [M+2]$^{2+}$=1068.1;<br>[M+3]$^{3+}$=712.3 |
| Cy-9B scrambled | Ac-LLLDAGAGAAGAGQ-$NH_2$ | [M+1]$^{+}$=1226.35;<br>[M+2]$^{2+}$=613.7;<br>[M+3]$^{3+}$=409.4 | [M+2]$^{2+}$=613.6;<br>[M+3]$^{3+}$=409.2 |
| Biot-scrambled | Biot-[$O_2$Oc]-LLLDAGAGAAGAGQ-$NH_2$ | [M+1]$^{+}$=1455.4;<br>[M+2]$^{2+}$=723.7;<br>[M+3]$^{3+}$=482.8 | [M+2]$^{2+}$=723.9;<br>[M+3]$^{3+}$=482.9 |
| Cy-9B-Cys Scrambled | Ac-LLLDAGAGAACAGQ-$NH_2$ | [M+1]+=1272.4;<br>[M+2]2+=636.7;<br>[M+3]3+=424.8 | [M+1]+=1272.5;<br>[M+2]2+=636.7;<br>[M+3]3+=424.9 |

**Table S1. List of peptides used in this study.** Sequences of the peptides synthesized with an acetylated (Ac) N-terminus and an amidated ($NH_2$) C-terminus. Molecular mass of synthetic peptides as determined by mass spectrometry analysis (ESI-MS). Standard amino acids are denoted using the one-letter code. Noncanonical residues are indicated as: X1 = (N3)K, azidolysine; X2 = Prg, propargylglycine.

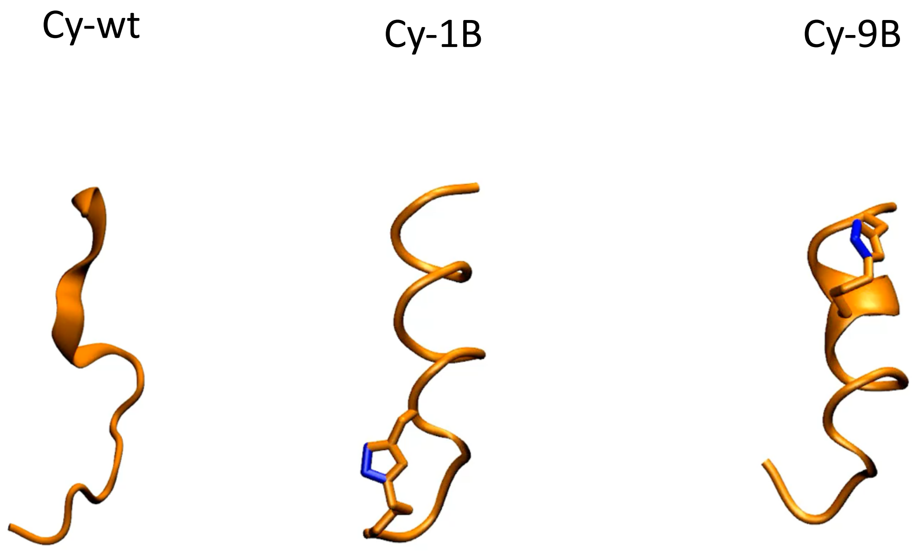


**Video S1. Molecular dynamics simulations of peptide conformations.** The three videos illustrate the structural behavior of the peptides over the course of the MD simulations, highlighting their ability to maintain alpha-helical folding: linear peptide (Cy-wt), N-terminal stapled peptide (Cy-1B), and C-terminal stapled peptide (Cy-9B).
